# A neural circuit model for human sensorimotor timing

**DOI:** 10.1101/712141

**Authors:** Seth W. Egger, Nhat M. Le, Mehrdad Jazayeri

## Abstract

Humans can rapidly and flexibly coordinate their movements with external stimuli. Theoretical considerations suggest that this flexibility can be understood in terms of how sensory responses reconfigure the neural circuits that control movements. However, because external stimuli can occur at unexpected times, it is unclear how the corresponding sensory inputs can be used to exert flexible control over the ongoing activity of recurrent neural circuits. Here, we tackle this problem in the domain of sensorimotor timing and develop a circuit-level model that provides insight into how the brain coordinates movement times with expected and unexpected temporal events. The model consists of two interacting modules, a motor planning module that controls movement times and a sensory anticipation module that anticipates external events. Both modules harbor a reservoir of latent dynamics and their interaction forms a control system whose output is adjusted adaptively to minimize timing errors. We show that the model’s output matches human behavior in a range of tasks including time interval production, periodic production, synchronization/continuation, and Bayesian time interval reproduction. These results demonstrate how recurrent interactions in a simple and modular neural circuit could create the dynamics needed to control temporal aspects of behavior.

## Introduction

A hallmark of human behavior is the ability to rapidly and flexibly coordinate movements with incoming sensory cues^1–7^. For example, a pianist can synchronize the tempo of his movements to beats of a music, and a tennis player can rapidly adjust her movements to return a serve. The remarkable flexibility with which we coordinate our movements with external events has been a major topic of research in psychophysics, systems neuroscience, and robotics. However, progress in developing models of sensorimotor coordination has been limited because we still lack a rigorous framework to understand how networks of neurons can flexibly and systematically adjust their behavior in response to inputs.

Recent studies have proposed that flexible sensorimotor coordination may be understood using the language of dynamical systems^8–11^. The key intuition is that recurrent networks of neurons establish versatile dynamical systems^12–16^ that can generate a diverse range of possible of outputs that qualitatively match the patterns of neural activity in the premotor and motor areas responsible for the control of movement dynamics^9, 17–20^. Within this framework, a sensory input to a recurrent neural network can be viewed as a control signal that reconfigures the system’s latent dynamics and allows the system to generate different outputs. However, recurrent neural network models are typically complex and do not offer the level of interpretability needed to engineer circuits that could use inputs for flexible sensorimotor coordination.

Here, we tackle this question within the domain of time, asking how a simple and interpretable neural circuit could flexibly coordinate the timing of its output to external temporal events. Timing provides a prime example of sensorimotor coordination and is crucial in behaviors that demand the generation of delayed motor responses, generation of rhythmic movements with a desired tempo, and synchronization of movements to anticipated events. Early experiments found that when animals are asked to delay their motor responses, many neurons exhibit ramping activity during the delay period; i.e., activity that increases or decreases monotonically over time^21–23^. These experiments inspired a simple ramp-to-threshold model for action initiation. According to this model, actions are initiated reliably when the ramping activity reaches a fixed threshold^24, 25^. When the threshold is fixed, flexible temporal control requires the system to adjust the rate of change, or speed, at which neural dynamics evolve. Indeed, it has been shown that neural circuits generate dynamics with a speed that has an inverse relationship to the instructed interval duration^9, 20, 26^.

While the adjustment of speed is a common empirical observation, the mechanisms by which this is achieved is not fully understood. Recently, it was shown that flexible control of speed can be achieved through nonlinear interactions within a simple recurrent neural network model^20^. This model relies on a simple circuit consisting of a pair of units with reciprocal inhibitory connections. By providing an appropriate shared tonic input to these two units, a ramp-to-threshold with the desired speed can be generated. (Figure 1a). From the perspective of control, this model can be viewed as an open-loop controller that converts an instruction (i.e., shared input) to the desired dynamics (i.e., ramp with the desired speed).

**Figure 1.**
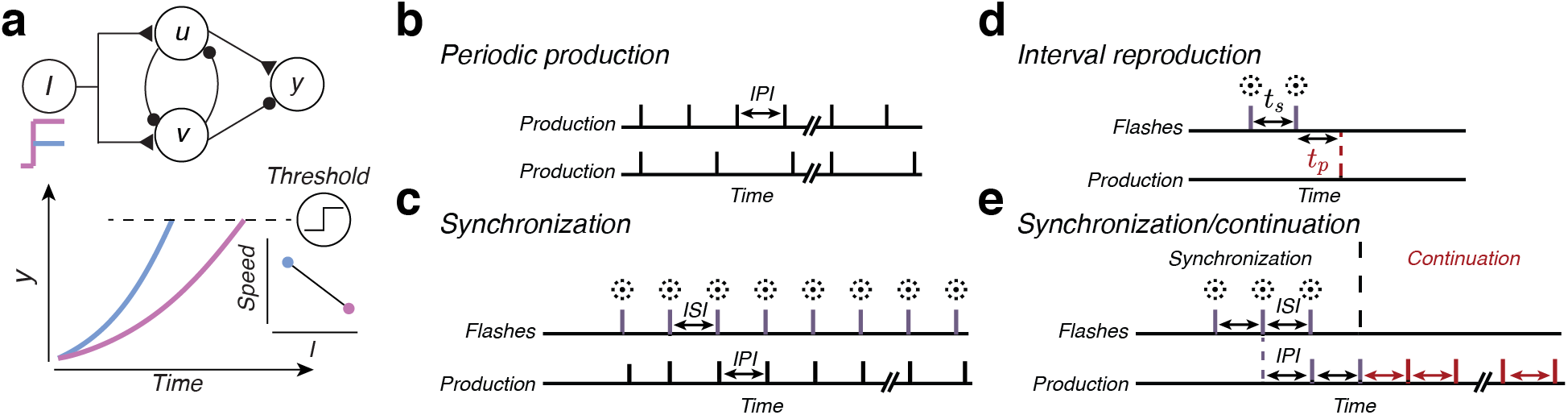
Basic circuit module for timed action and repertoire of human timing behavior. a) Basic circuit module with excitatory connections indicated by triangles and inhibitory connections indicated by circles (top). Two units, *u* and *v*, reciprocally inhibit each other. Under input from a unit, *I*, the activity of *u* and *v* will evolve dynamically to a state where *u* or *v* dominates. The state along this dynamic trajectory is read-off by a unit, *y*, which receives excitatory input from *u* and inhibitory input from *v*. Setting the initial conditions such that *u* dominates leads to an increase in activity of *y* over time (bottom) until its level of activity reaches a threshold for action initiation (dashed line). Colors indicate the output of *y* under large (pink) or small (blue) input, and the inset indicates the relationship between the level of input and the speed at which neural activity evolves. (b-e) Classic timing tasks used to study human timing behavior b) Periodic production requires the subject to produce a series of actions over time (vertical lines), with a constant inter-production interval (IPI). Top and bottom are examples of two different IPIs. c) Synchronization requires the subject to time a series of actions (vertical black lines) such that they are simultaneous with a series of sensory inputs (e.g. flashes; vertical magenta lines) with a set interstimulus interval (ISI). d) Interval reproduction requires the subject to measure an interval, *t*_*s*_, sampled from a set distribution of possible intervals, and to produce an interval, *t*_*p*_, that matches *t*_*s*_ as accurately as possible. e) Synchronization/continuation requires the subject to synchronize actions to a series of inputs with an ISI selected at random from a prior distribution and then, after the stimulus is extinguished, continue to produce actions with an IPI matching the ISI selected on a given trial.

However, the larger utility of this model depends on whether it can be extended to accommodate additional constraints associated with temporal control of movements. In particular, temporal control of movements has to accommodate internal noise within the nervous system^27^, uncertainty due to unanticipated changes in the environment^28^, and sensorimotor delays imposed by internal and external temporal contingencies. These factors limit the capacity of open-loop systems to achieve robust temporal control. Efficient control systems often rely on sensory feedback to combat noise, and when sensory feedback is delayed, they rely on a mechanism to predict future sensory and motor states^29–31^. Here, we augment the original open-loop system by adding two critical components, a sensory anticipation module and a feedback mechanism. We then verify that the augmented model can accommodate internal noise, external uncertainty, and large sensorimotor delays, and demonstrate that it captures key features of human behavior in a number of classic timing tasks (Figure 1b-e).

## Results

We will describe the full model in four steps. We start by introducing a basic circuit module (BCM) that acts as a flexible open-loop controller for producing desired time intervals. We extend the BCM to a motor planning module (MPM) capable of producing isochronous rhythms. We then introduce a sensory anticipation module (SAM) that provides the means for anticipating and predicting upcoming temporal events. Finally, we introduce the full model that combines the MPM and SAM to create a system that can dynamically coordinate motor plans and actions with anticipated and unanticipated stimuli in a range of behavioral tasks.

### Basic circuit module (BCM) for interval production

A recent study proposed a simple model of flexible action timing that leverages nonlinear dynamics to generate ramp-like activity which leads to a motor output after this activity crosses a fixed threshold^20^. In this circuit, which we refer to as the basic circuit module (BCM), the slope of the ramp-like activity can be adjusted by an instructive input such that the threshold is crossed after a desired delay. This is achieved mechanistically through two units, *u* and *v*, which represent two populations of neurons with recurrent inhibitory connections (Figure 1a). Each unit’s output represents the average activity of the population. When driven by a shared, excitatory tonic input, *I*, the activity of the system evolves towards one of two fixed points, with *u* dominating or *v* dominating, depending on the initial conditions. Importantly, the nonlinear input/output functions of *u* and *v* result in an interaction between the level of *I* and the energy landscape of the BCM, allowing the input to control the speed at which the activity of *u* and *v* evolves^20^. *u* and *v* were modeled as rate units according to

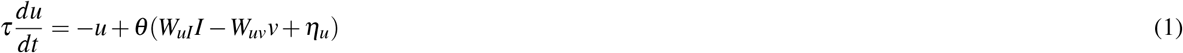

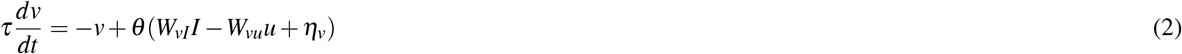

*W*_*uI*_ and *W*_*vI*_ denote the weight of an excitatory input, *I*, driving units *u* and *v*, respectively. *W*_*uv*_ and *W*_*vu*_ denote the strength of inhibitory coupling from *v* to *u*, and from *u* to *v*, respectively. *τ* is the time-constant of each unit, which was set to *τ* = 100 ms for all units, a value consistent with previous models^20, 32, 33^. *η*_*u*_ and *η*_*v*_ are stochastic inputs to each unit and are modeled as independent white noise with standard deviation *σ*_*n*_. *θ* is the nonlinear function with outputs between zero and one that specifies the output rate of *u* or *v* as a function of its synaptic inputs. The critical feature for speed control is that *θ* is a sigmoidal function. Here, we choose *θ* (*x*) = 1*/*[1 + exp(− *x*)]^20^. In subsequent modeling, we assume *W*_*uI*_ = *W*_*vI*_ (identical shared excitatory input) and *W*_*uv*_ = *W*_*vu*_ (symmetric mutual inhibition).

A straightforward neural circuit for action timing can be constructed by comparing the level of activity of *u* and *v* according to a simple differencing mechanism implemented by a linear unit, *y*, that receives excitatory input from *u* and inhibitory input from *v*. The dynamics of *y* were modeled as

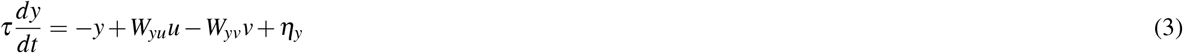

where *W*_*yu*_ and *W*_*yv*_ specify the strength of coupling from *u* to *y* and *v* to *y*, respectively. Here we take *W*_*yu*_ = *W*_*yv*_ = 1. As in the units *u* and *v*, we assume *y* is subject to independent, stochastic input *η*_*y*_ with standard deviation *σ*_*n*_. This noise term, combined with the noise injected into *u* to *v*, generates variability in threshold crossing times.

To illustrate the behavior of the BCM, we initialized the module such that *u* dominates *v*, and characterized the dynamics of the output, *y*, for different levels of the shared input. As expected, driving the BCM with a shared tonic input, *I*, causes the output to exhibit a nonlinear ramping activity (Figure 1a) whose rate of increase is inversely related to *I*. Accordingly, the BCM can readily produce different time intervals if we assume that movement initiation is triggered when *y* reaches a fixed threshold, *y*_0_. Therefore, the BCM converts a time interval instruction conveyed by a tonic input to a dynamic output that evolves to threshold at an appropriate speed to achieve the desired produced interval (Figure 1a). In other words, the BCM acts as an open-loop controller for producing a single instructed time interval^20^.

### A reset mechanism for periodic production

We next considered the production of multiple movements with the interproduction interval (IPI) set according to an instruction signal. This is challenging with the basic module because the nonlinear dynamics of the BCM dictate that *u* and *v* evolve to a fixed point^20^, limiting its output to a single action. One solution to this problem is to utilize a series of concatenated BCMs, and use threshold crossing in each module to trigger the dynamics of the next. However, this is a highly unrealistic solution as it requires the number of modules to grow with the number of actions.

An alternative strategy is to use a single module repeatedly via a mechanism that can reset *u* and *v* after each threshold crossing, allowing the model to generate arbitrary number of timed outputs. To implement this solution, we augmented the BCM such that every threshold crossing would additionally serve as an internal reset signal. From a neurobiological perspective, this internal signal can be viewed as a corollary discharge that is activated concurrently with action initiation^34–36^. This modified architecture, which we call the motor planning module (MPM), consists of four units, *u*_*p*_, *v*_*p*_, *I*, and *y*_*p*_. As in the BCM, the input unit *I* drives dynamics in the MPM circuit such that *y*_*p*_ evolves toward a fixed threshold, *y*_0_, for motor initiation. The additional corollary discharge is modeled as a transient, 10 ms pulse, *I*_*p*_, that is activated right after the time of threshold crossing (Figure 2a, dashed line). Mathematically, this can be formalized by augmenting equations 1 and 2 with the input *I*_*p*_ according to

**Figure 2.**
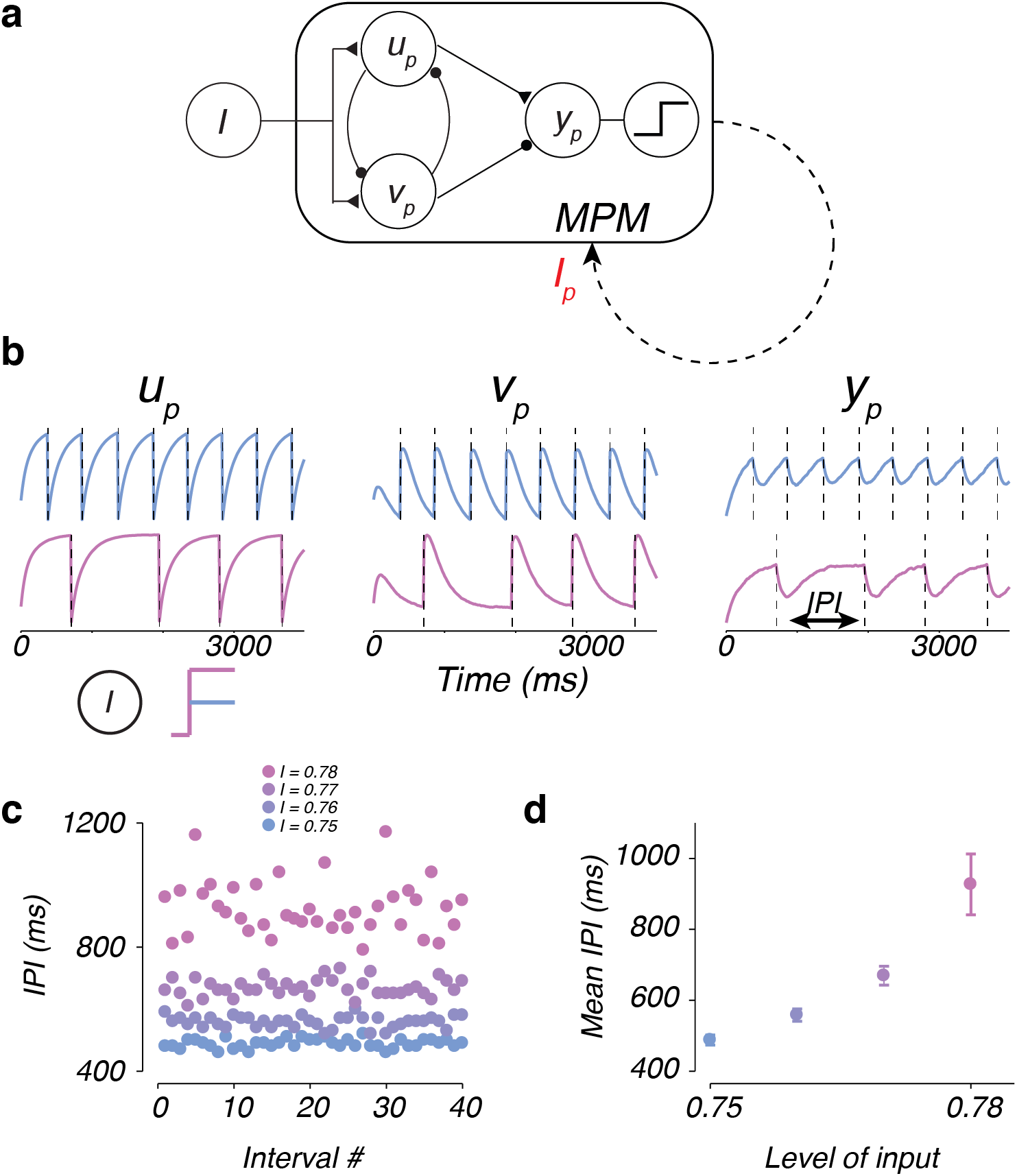
Periodic production of different interval durations is achieved by adjusting the level of input to the motor planning module (MPM) (a) The architecture of the MPM is identical to the BCM, with the addition of an action initiation trigger once *y*_*p*_ crosses threshold (represented by the nonlinear unit at right). The action initiation signal is then fed back to the MPM as an input, *I*_*p*_ (dashed line). Excitatory connections are indicated by triangles and inhibitory connections are indicated by circles. (b) Behavior of the units *u*_*p*_, *v*_*p*_ and *y*_*p*_ in response to a small (blue) or large (pink) input, *I*. Vertical dashed lines indicate the timing of threshold crossing and motor output. The time between successive motor outputs is defined as the inter-production interval (IPI). (c) Value of the first 40 IPIs generated by the MPM at each level of input, *I* (colors), in representative trials. (d) Mean IPI (*±* standard deviation) as a function of input, *I*.

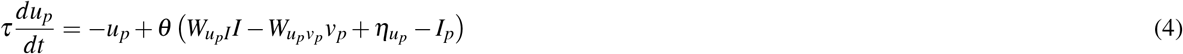

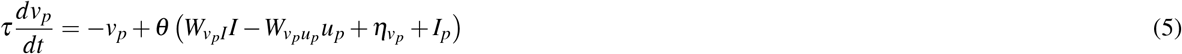

We can gain some intuition for how the MPM functions by examining traces of the activity of *u*_*p*_, *v*_*p*_, and *y*_*p*_ over time when *I* is small or large (Figure 2b, blue and pink, respectively). In both contexts, the behavior of the units is similar. From the onset of the trial, the activity of *u*_*p*_ increases and *v*_*p*_ decreases, resulting in *y*_*p*_ increasing over time. Once *y*_*p*_ reaches *y*_0_, the module generates an action (vertical dashed lines), *I*_*p*_ resets *u*_*p*_ and *v*_*p*_, and dynamics begin to unfold again. Because *y*_*p*_ is driven by *u*_*p*_ and *v*_*p*_, this results in a rapid drop in the output of *y*_*p*_ before starting another ramp to threshold. Importantly, because the MPM is identical to the BCM after each reset, we expected that the speed at which *y*_*p*_ ramps up over time would depend on the level of *I* (Figure 2b). Simulations of the MPM verified this prediction: IPI was progressively longer for larger *I* (Figure 2c), and across multiple simulations, IPI and *I* were strongly correlated (Figure 2d; *r*^2^ = 0.84; *F*(1, 160) = 828.6; *p* ≪ 0.01). Moreover, in the presence of noise injected into the circuit (*σ*_*n*_ = 0.01, see Methods), larger *I* increased the variability of IPI (one-tailed F test; *I* = 0.75 to *I* = 0.76: *p* = 0.014*, F*(80, 70) = 0.60; *I* = 0.76 to *I* = 0.77: *p <* 0.01*, F*(57, 70) = 2.95; *I* = 0.77 to *I* = 0.78: *p <* 0.01*, F*(40, 57) = 10.93). In other words, IPI variability was larger for longer IPI, which is a hallmark of isochronous rhythm production in humans^37^.

### A sensory anticipation circuit for predicting future temporal events

The MPM establishes a circuit for producing isochronous rhythms for which IPI can be adjusted flexibly by the input level. However, in many natural circumstances, the IPI is not known in advance and has to be adjusted based on external temporal events^7, 28, 38^. For example, a guitarist who wants to synchronize actions with a drummer may not know the correct tempo before hearing the drummer play a few beats. For the MPM to emulate this type of behavior, the input to the MPM must be adjusted dynamically depending on the interval between temporal events. In general, measuring time between events can be achieved in two ways. One way is to implement a circuit whose output is a ramp with a fixed slope. In such a circuit, the output level would provide a moment-by-moment estimate of elapsed time (Supplementary Figure 1a). Alternatively, the circuit may function predictively, and adjust the slope of the ramp such that the output would reach a certain expected level at the anticipated time of the next event (Supplementary Figure 1b).

Physiological recordings in the frontal cortex suggest that the nervous system relies on the latter predictive mechanism^8, 39^. Specifically, it was found that when monkeys anticipate a stimulus later in time, neural signals in the frontal cortex evolve more slowly. Conversely, when monkeys anticipate a stimulus earlier in time, neural signals evolve more quickly. As a result, population activity reaches the same terminal state when the sensory event is expected. This mechanism is remarkably similar to how the MPM functions, which is to adjust the slope of a ramp so that the output would reach a threshold at a desired time. Therefore, we leveraged the computational logic of MPM to create a sensory anticipation module (SAM) that could predict the time of upcoming sensory events (Figure 3a). Like the MPM, the SAM receives an input, *I*, that controls the speed of dynamics. However, unlike the MPM, the output unit of the SAM, *y*_*s*_, does not generate an action when it crosses the threshold *y*_0_. Instead, *y*_*s*_ indicates the output relative to *y*_0_ at the time of sensory feedback. Therefore, on average, a value of *y*_*s*_ < *y*_0_ indicates a slower than needed tempo. Accordingly, *I* should be decremented to speed up system dynamics such that *y*_*s*_ approaches *y*_0_ at the correct time. Conversely, if *y_s_ > y*_0_, *I* should be incremented to produce the correct timing. In other words, to generate the appropriate control output, the input must be adjusted such that the SAM generates dynamics that match *y*_*s*_ to *y*_0_ at the time of the next stimulus.

**Figure 3.**
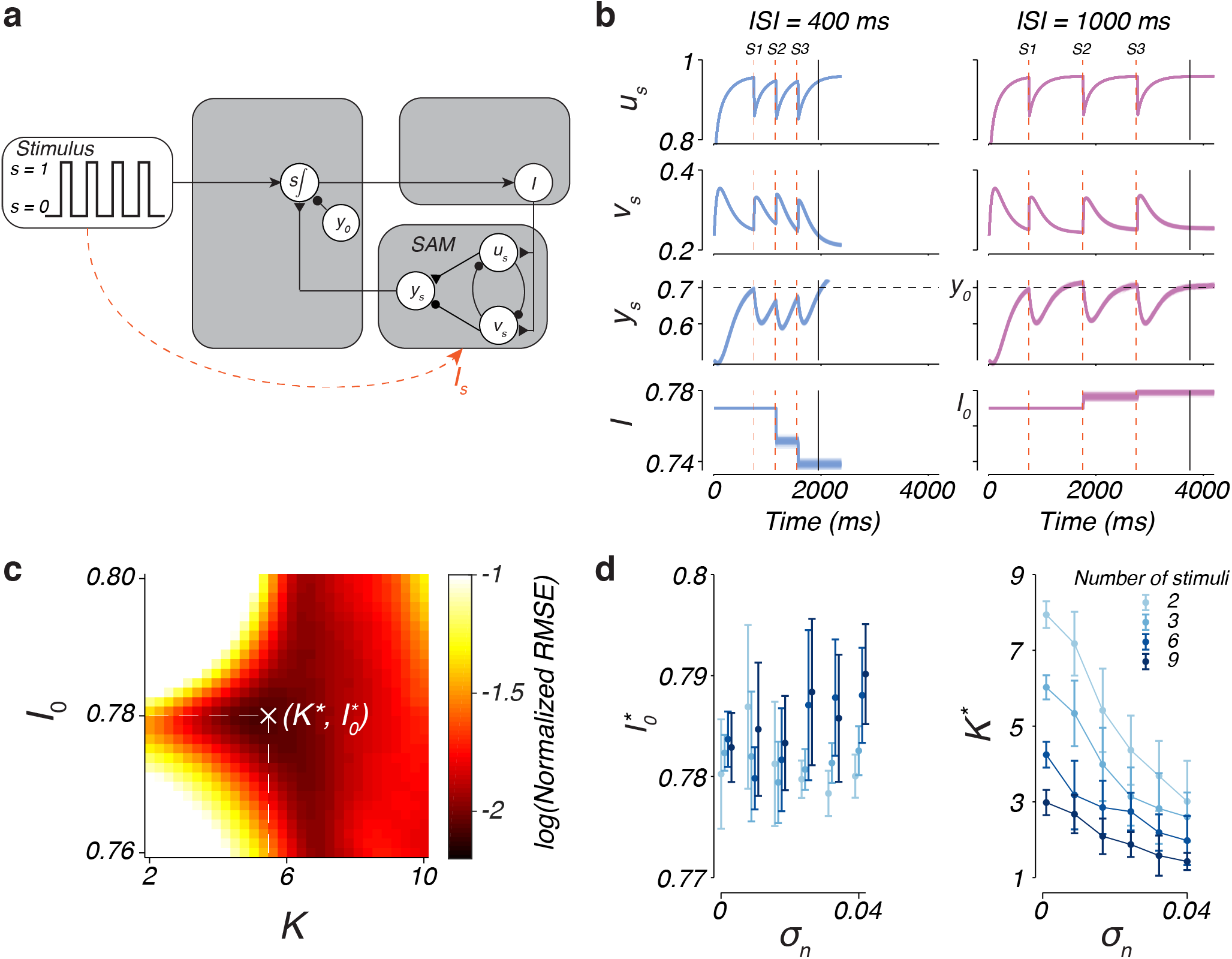
Dynamic adjustment of input with the sensory anticipation module (SAM). a) Architecture of the updating mechanism and SAM. The updating mechanism, represented by s∫, relies on difference between the output of the SAM, *y*_*s*_, and the desired level of activity at the time of each stimulus. This is implemented by integrating the summed activity of *y*_*s*_ and a tonically firing inhibitory unit, *y*_0_, into the input unit, *I*, when the stimulus is on (*s* = 1). When the stimulus is off (*s* = 0), integration is prevented and *I* remains constant. Dashed line indicates a stimulus input, *I*_*s*_, directly to *u*_*s*_ and *v*_*s*_ when *s* = 1, resetting the SAM after each stimulus. Excitatory connections are indicated by triangles, inhibitory connections are indicated by circles, and non-specific synapses are indicated by arrows. b) Response of the SAM units *u_s_, v_s_, y*_*s*_, and *I* to three isochronous stimuli (S1, S2, S3, vertical dashed lines) with an ISI of 400 ms (blue) and 1000 ms (pink). The vertical black line indicates the time of the stimulus which is to be predicted. Each trace represents the response of a unit in one of 100 different trials. The horizontal dashed lines indicate the threshold *y*_0_. c) Example optimization of the value of the weight given to the update, *K*, and *I*_0_, at noise level *σ*_*n*_ = 0.005 and *N* = 3 stimuli. Color scale represents the RMSE for each pair of *K* and *I*_0_ tested. d) The optimized parameter values, *K*^*∗*^ and 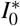, as a function of the level of the level of noise, *σ*_*n*_, and number of input stimuli (colors). Error bars indicate the standard deviation across optimization runs.

To implement the adjustments to *I* specified by the above logic, we set the change in *I* over time to a rate proportional to the error signal, *y*_*s*_ − *y*_0_, according to

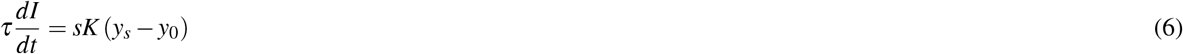

This update rule achieves our original goal because when 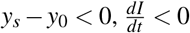, resulting in a decrease in *I*. Conversely, when 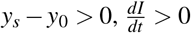 and *I* will increase. The rate of change of *I* is related to the error signal *y*_*s*_ − *y*_0_ through a constant *K*, which is a free parameter of the model. Furthermore, to enforce this update only during the time of sensory feedback, we make use of a variable, *s*, which indicates whether the sensory feedback is on or off. Therefore, when feedback is on (*s* = 1), *I* is adjusted, but when there is no feedback (*s* = 0), 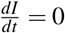, and *I* is held constant.

Figure 3b illustrates the response of SAM units to a series of three isochronous sensory inputs (S1, S2, and S3; vertical dashed lines) with a short (400 ms) or long (1000 ms) ISI. We initialize the module by setting the input, *I*, at *I*_0_ and allowing dynamics to run forward for 750 ms before providing the S1 stimulus (see Methods). The S1 stimulus acts to initialize dynamics by providing a transient input, *I*_*s*_, that resets *u*_*s*_ and *v*_*s*_ in a manner analogous to that used to reset the MPM (Figure 3a; see Methods). Following the transient input, dynamics of the SAM will unfold such that *y*_*s*_ increases over time. For a short ISI, the value of *y*_*s*_ at the time of the S2 stimulus is lower than *y*_0_, indicating that the module needs to be sped up to appropriately time the response (Figure 3b, left). In contrast, for a long ISI, the value of *y*_*s*_ is greater than *y*_0_, indicating that the dynamics need to be slowed (Figure 3b, right). Following the update, the S2 stimulus again resets *u*_*s*_ and *v*_*s*_ by providing the transient input, *I*_*s*_, allowing the dynamics of the SAM to proceed again driven by an adjusted value of *I* (Figure 3b, bottom). By adjusting *I* in this manner, the response of *y*_*s*_ at the time of the S3 stimulus is closer to *y*_0_. Iterating the updating process after each stimulus input of an isochronous sequence, the SAM can dynamically adjust its output such that *y*_*s*_ will eventually match *y*_0_ at the precise time of each stimulus input.

In effect, adjusting the input to the SAM such that *y*_*s*_ matches *y*_0_ at the time of feedback renders the dynamics predictive of the stimulus timing. Therefore, the SAM provides a plausible mechanism to implement prediction-based estimation algorithms hypothesized to underlie human timing behavior in a variety of tasks^40^. However, the prediction performance of the SAM depends on the interaction between the noise (*σ*_*n*_), initial input (*I*_0_), update constant (*K*), and the number of input stimuli (*N*) that the system receives. We therefore explored the impact of these parameters on the circuit’s ability to predict the time of the *N*^th^ stimulus of a sequence (e.g. S4 in Figure 3b, vertical solid line).

To quantify the predictive ability of the SAM, we measured the root-mean-squared-error (RMSE) between the predicted time (the time at which *y*_*s*_ crosses *y*_0_) and the time that the (*N* + 1)^th^ input would have occurred, across 100 simulated trials in which the ISI was sampled from a uniform prior between 600 and 1000 ms. We then repeated this process for different combinations of *K* and *I*_0_ to determine the impact of these parameters on the prediction performance. Figure 3c shows how the RMSE varies as a function of *K* and *I*_0_ when *N* = 3 and *σ*_*n*_ = 0.005. For values of *I*_0_ that are large and *K* that are small, the circuit generates large errors in timing. In contrast, intermediate values of *I*_0_ and *K* lead to the smallest errors in timing, and a specific combination of 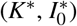 minimizes the RMSE (see Figure 3c). Exploration of the parameter combinations that minimize errors for different levels of noise and stimulus number *N* revealed three key results. First, 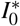 is largely independent of either the level of noise or the number of stimuli presented (Figure 3d, left). This is consistent with *I*_0_ acting as a control signal that reflects an initial estimate of the ISI based on expectations before any feedback is integrated. Second, *K*^*∗*^ decreases with increasing *σ*_*n*_ (Figure 3d, right), indicating that as internal noise increases, the error signals generated by the SAM are less reliable and therefore should have a smaller influence on the update mechanism. Third, as *N* increases, *K*^*∗*^ decreases, indicating that the weighting given to each error is decreased in favor of integrating across feedbacks (Figure 3d, right).

### Sensorimotor updating by combining the SAM and MPM

So far, we found that the MPM can produce different IPIs depending on the level of input, and the SAM can adjust the input level based on the anticipated time of sensory events. Accordingly, we reasoned that the SAM and MPM might together be able to generate timed outputs that are in register with incoming sensory events: the SAM would use its predictive mechanism to adjust the input based on sensory events and the MPM would use that input to adjust IPI. We therefore connected the SAM to the MPM by having them share the input, *I*, (Figure 4a) and then tested the ability of the MPM to match its IPI to a stimulus with an ISI that changes in a blocked fashion (every 20 beats) over the course of a trial (Figure 4b).

**Figure 4.**
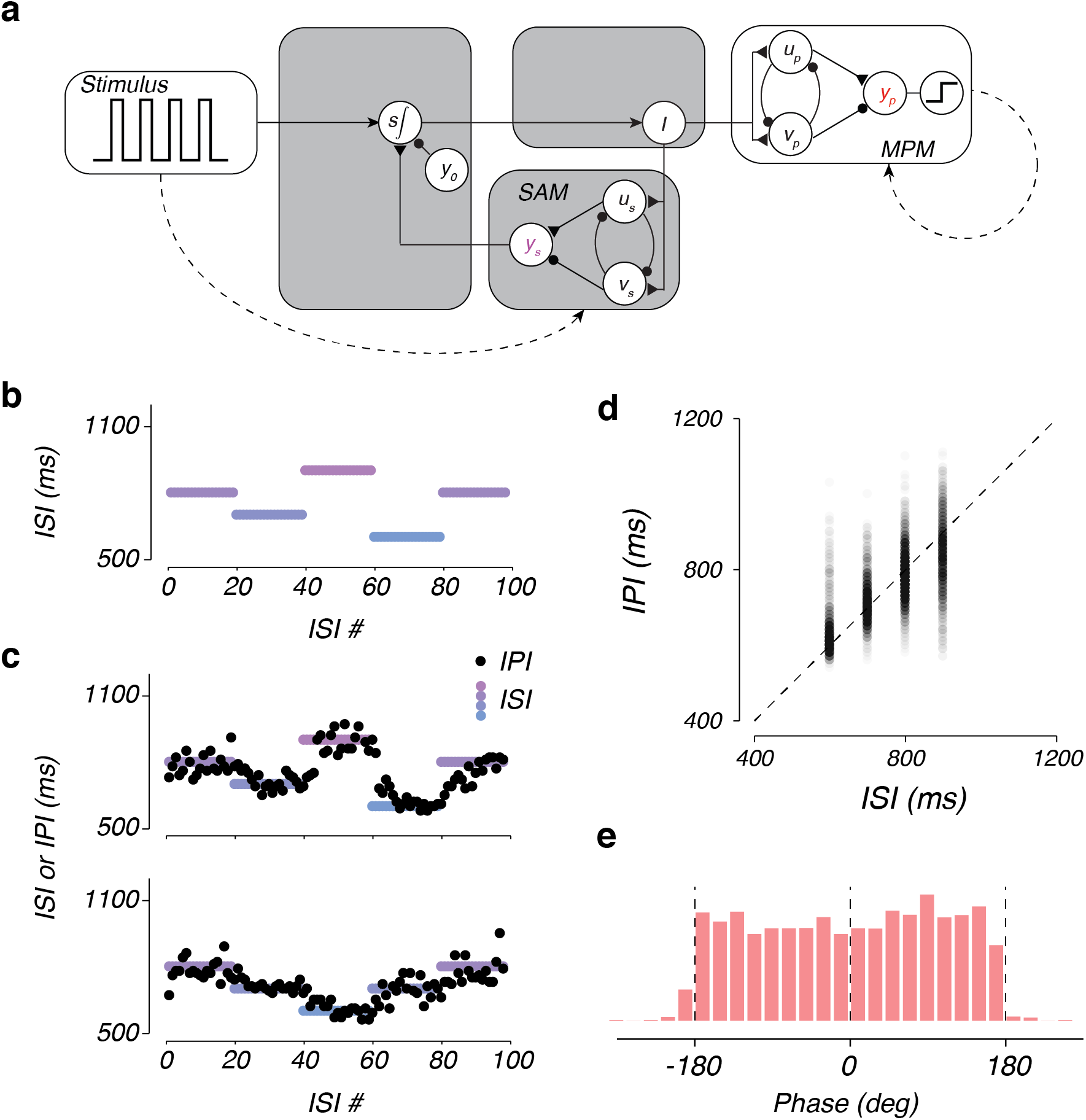
Online adjustment of inputs allows a combined circuit to match IPI to stimulus ISI. a) Wiring diagram of the circuit model that combines the sensory anticipation module (SAM) with the motor planning module (MPM). Conventions as in Figures 2 and 3. b) Example inter-stimulus-interval (ISI) sequence for a trial of the ISI tracking task. Colors indicate different ISIs. We initiated each trial with a block of twenty consecutive 800 ms ISIs. Each subsequent block of 20 ISIs was randomly selected from a discrete uniform distribution between 600 and 900 ms. Each trial consisted of 5 blocks and 100 total ISIs. c) Inter-production intervals (IPIs; black circles) associated with each ISI (colors) for two example trials of the ISI tracking task. d) Density of IPIs (grayscale) as a function of the associated ISI. Dashed line plots indicates perfect tracking. e) Distribution of the phase of motor output. A phase of zero indicates output that is synchronous with the stimulus input.

As expected, the circuit was able to successfully track ISI throughout the run (Figure 4c; see Methods). After each block transition, the circuit gradually adjusts its output to match the new randomly chosen ISI. After each transition, the discrepancy between the expected and actual ISI causes the SAM to generate a prediction error. This error adjusts the level of *I* according to the update constant *K* (see Methods), and changes the speed of dynamics in both modules such that the next IPI would, on average, more closely match the new ISI. With each subsequent ISI, the SAM further adjusts *I* until the average IPI matches the ISI. This process is repeated for every change in ISI, allowing the circuit to successfully track the ISI with corresponding changes to the IPI (Figure 4c). Across trials and ISI, the circuit successfully generates IPIs to match the stimulus ISI (*r*^2^ = 0.53; *p* ≪ 0.01; Figure 4d), demonstrating that a circuit comprised of the SAM and MPM can flexibly adjust its timed motor output in accordance with discrete sensory inputs.

### Full circuit model for synchronization

Coupling between the MPM and SAM allowed the circuit to adaptively match the frequency of its output as indexed by the IPI to the frequency of input specified by the ISI. However, an important related question is whether the circuit’s output matched the input in phase in addition to frequency. This is an important consideration as humans can readily produce rhythmic actions that are synchronized (phase-matched) to external beats^41^. Therefore, we examined the difference in absolute timing, or asynchrony, between each stimulus input and motor output of the circuit performing the ISI tracking task. Across trials and ISIs, the distribution of phase differences between sensory inputs and motor outputs produced by the circuit was uniformly distributed between −180° and 180° (Figure 4e). This indicates that, although the circuit correctly matched the IPI to the ISI, the timing of motor outputs had no systematic relationship to the timing of the input stimuli.

The source of this problem is that the circuit has no mechanism to determine whether the MPM output leads or lags the stimulus. Intuitively, this problem may be addressed by a separate system that measures the time difference between motor outputs and stimulus inputs^42–44^. Adjustments to phase can then be achieved by adjusting the desired timing of the next output.

Implementing this using neural circuits is challenging, however, because the clock which measures this difference must start at either the time of the stimulus or motor output, depending on whether the motor out lags or leads the stimulus.

Previous circuit models have address this through highly nonlinear phase adjustment functions^45–47^. Our circuit, however, provides a simple alternative solution. Since the ramping output in the SAM (*y*_*s*_) and MPM (*y*_*p*_) are synchronized to sensory and motor events, respectively, the difference between *y*_*s*_ and *y*_*p*_ provides an analog signal reflecting their relative phase. Specifically, when *y_s_ > y*_*p*_, the MPM is lagging the stimulus and therefore should be sped up by decreasing the total input to the MPM. Similarly, when *y_s_ < y*_*p*_, the MPM is leading the stimulus and, therefore, should be slowed down by increasing the total input to the MPM. To implement this solution, we augment the input *I* with a signal ∆*I* of the form

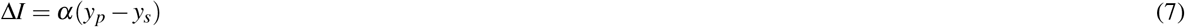

where *α* determines the weight given to the difference signal. This adjustment can be realized by a unit which receives excitatory and inhibitory input form *y*_*p*_ and *y*_*s*_, respectively. (Figure 5a, cyan). By using the sum of ∆*I* and *I* to drive the MPM, the dynamics of the MPM could be slowed or sped up according to its phase relative to the stimulus. Figure 5b uses an example to demonstrate this correction scheme. Initially, *y*_*p*_ slightly lags *y*_*s*_. This asynchrony generates a biphasic ∆*I* whose value is transiently positive (between the stimulus onset and motor output), and then negative until the next stimulus onset (vertical dashed lines). The addition of this error signal reduces the total tonic drive to the MPM and increases the speed of its dynamics relative to the SAM. After this adjustment is applied over several ISIs/IPIs, the MPM becomes increasingly synchronized with the stimulus, and the error between *y*_*p*_ and *y*_*s*_ decreases. Implementing this strategy allowed the circuit to gradually reduce phase asynchrony and variability across intervals and ISIs (*F*(4899, 4899) = 2.25, *p* ≪ 0.01; Figure 5c). Intriguingly, the mean phase of motor output relative to the stimulus was slightly negative (−27.14° ± 71.45°), indicating the MPM slightly led the stimulus in a manner similar to humans performing analogous tasks^48–50^.

To examine the robustness of the model’s behavior with respect to the parameters, we examined the phase asynchrony for different values of *α* and *K*. Larger values of *α* makes the model more sensitive to phase errors between output and stimulus and allows the model to cancel phase asynchronies more effectively (Figure 5d, left panel). However, depending on task demands, this may not be ideal as larger values of *α* cause rapid beat-by-beat changes in IPI, which lead to reduced rhythmicity in the output and an overall increase of IPI error relative to ISI (Figure 5d, middle panel). Therefore, the optimal value of *α* depends on the relative cost of minimizing phase asynchrony and IPI error. For example, values between 0.1 and 0.15 allow the model to minimize the sum squared error of phase and IPI (Figure 5d, right panel). We also examined the effect of *K* on the model’s behavior and found that it has little effect on how *α* influences phase asynchrony and IPI error (Figure 5d, different colors).

**Figure 5.**
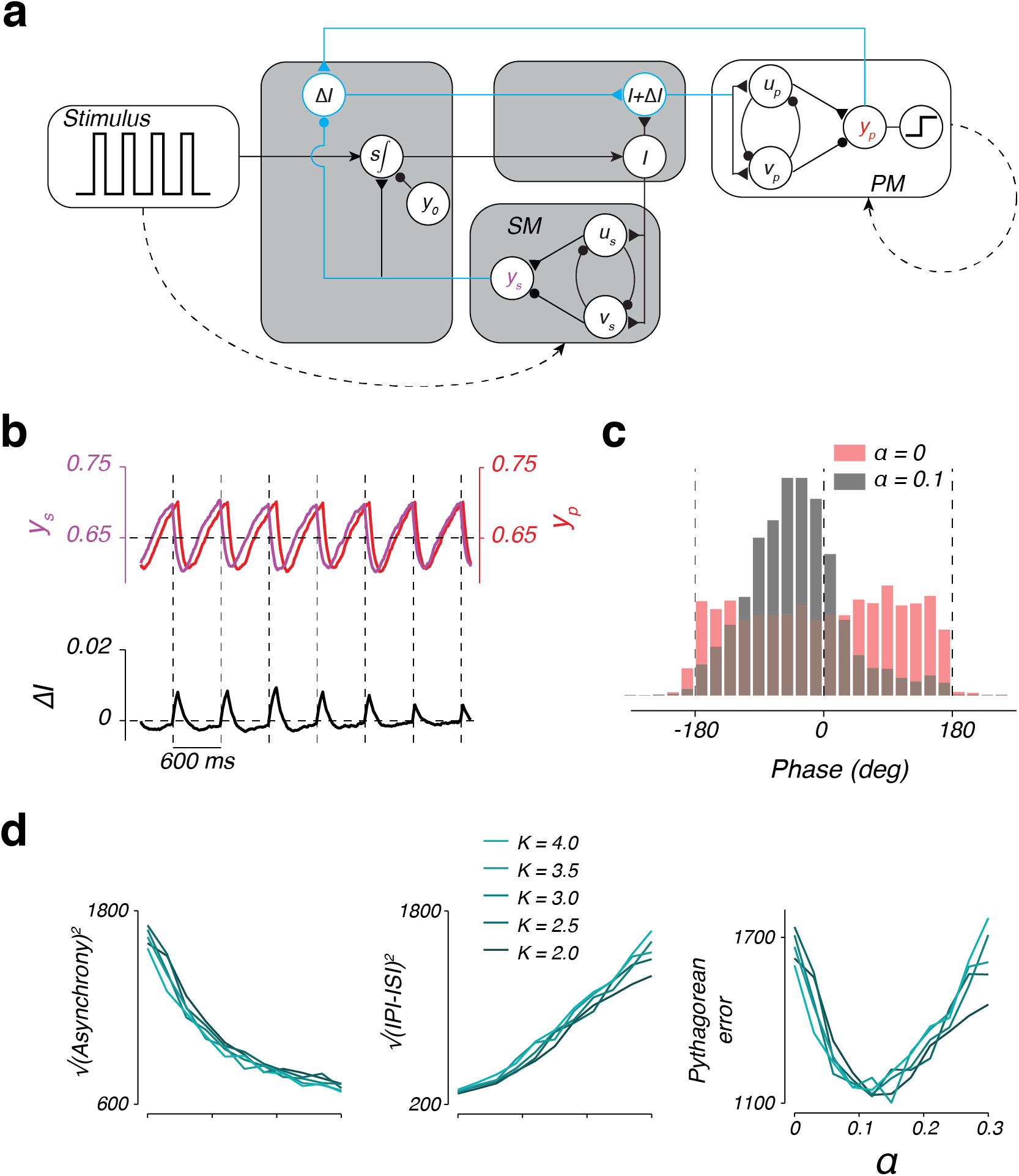
Full circuit model for synchronization. a) Augmented circuit model. To synchronize the MPM with SAM output, a second circuit pathway was introduced to measure the difference between these two outputs. The difference, weighted by *α*, augments the input to the plant with input ∆*I*. b) Plant output, *y*_*p*_, simulator output, *y*_*s*_, and their difference weighted by *α* = 0.1 (i.e. ∆*I* = *α*∆*y*). Augmenting *I* with ∆*I* adjusts controller output to the plant such that the time of production tends to match the time of flashes (vertical dashed lines). c) Distribution of the phase of production for the circuit with (black) and without (red) augmented input. d) Optimization of *α*. Increasing *α* decreases asynchrony (left). In contrast, increasing *α* increases the sum of the squared IPI errors (middle). As a result, a limited range of *α* values minimize Pythagorean errors, defined as 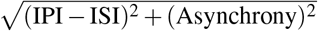 (right).

### Model’s behavior during stimulus perturbation is similar to humans

We compared the behavior of the model to that of humans in synchronization tasks during which the rhythmic input was perturbed in three different ways: a step change in ISI (Figure 6a, top), a phase shift (Figure 6b, top), and jittering a single event without changing the ISI (Figure 6c, top).

**Figure 6.**
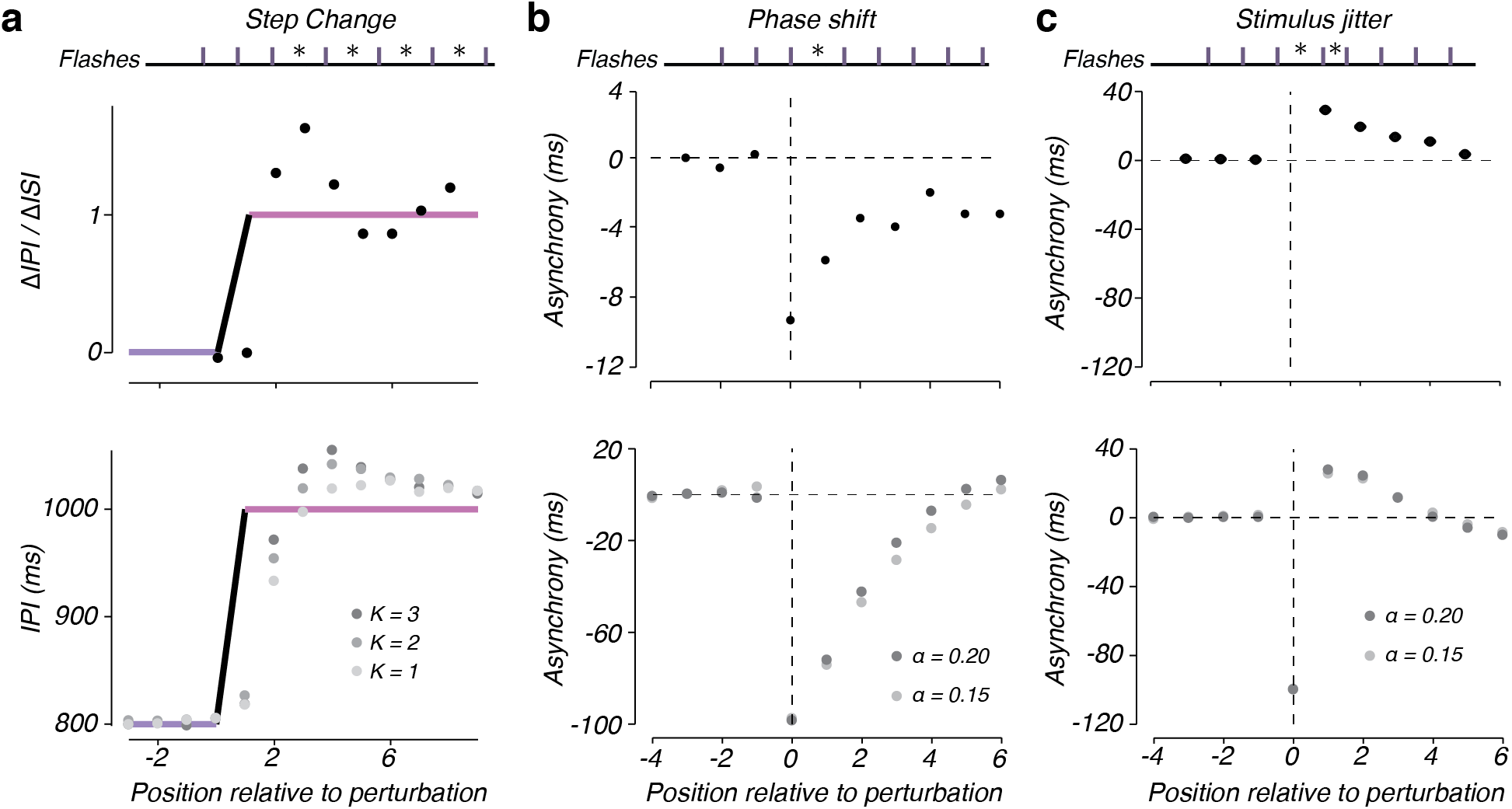
Circuit response to ISI perturbations. a) Model and human responses to a step change in the interstimulus interval (ISI). Top: schematic of the task design. Vertical bars indicate the timing of a sequence of flashes with the distance between vertical bars corresponding to the ISI. Asterisks denote ISIs that are perturbed relative to the initial ISI. Middle: example subject step change response reproduced from Michon, 1967^52^. Circles plot the change in interproduction interval (∆ IPI) relative to the change in ISI (∆ ISI). Bottom: average IPI of the circuit model (circles) to a step change in the ISI from 800 ms (thick blue line) to 1000 ms (thick pink line). The level of gray in the circles indicates circuit model behavior for different values of the parameter, *K*. b) Model and human responses to a phase shift in the stimulus timing. Top: schematic of the task design, as in panel a. A phase shift was induced by increasing the duration of one interval (asterisk). Middle: Average asynchrony measured in humans following a phase shift of 10 ms from a 500 ms ISI. Data reproduced from Repp, 2000^55^. Bottom: average asynchrony between the stimulus and the time of production of the circuit model (circles) following a phase shift of 100 ms from an ISI of 500 ms. To facilitate the comparison of the model to human behavior in response to the perturbation, we removed the mean asynchrony adopted before the perturbation from the mean responses of the circuit model. The horizontal dashed line indicates zero asynchrony and the vertical dashed line indicates the position of the first flash following the perturbation. The level of gray in the circles indicates circuit model behavior for different levels of the parameter, *α*. c) Model and human responses to temporal jitter of a single stimulus. Top: schematic of the task design, as in panel a. The timing of a single stimulus was jittered by increasing one interval by 100 ms, followed by decreasing the subsequent interval by 100 ms (asterisks) from an ISI of 500 ms. Middle: average asynchrony measured in humans following stimulus jitter. Data reproduced from Repp, 2002^58^ (asynchrony to the perturbed stimulus not shown). Bottom: average asynchrony between the stimulus and the time of production. Conventions as in panel b.

When facing a sudden change of ISI (Figure 6a, top), response times have to be adjusted in two ways. First, one has to compensate for the phase asynchrony caused by the sudden mismatch between action and stimulus. Second, one has to use the subsequent ISIs to appropriately adjust the IPI. For example, after an uncued increase in ISI, the first motor response would lead the stimulus. To regain synchrony, one has to delay the response to accommodate this phase lead, and additionally increase the subsequent IPIs to match the new ISI. In psychophysical experiments in humans, it has been shown that subjects adjust their response times by concurrently reducing asynchrony and adjusting IPI such that, after a few samples, actions are in sync with sensory inputs^43, 44, 50, 51^. A hallmark of this concurrent error correcting strategy is that IPIs exhibit a transient overshoot relative to the new ISI (Figure 6a, middle)^52–54^.

To test the behavior of the model in response to a change in ISI, we used the following simulation protocol: we allowed the model to reach steady-state for an ISI of 800 ms, stepped the ISI to 1000 ms, and measured subsequent IPIs produced by the model. We repeated this procedure 1000 times and measured how average IPI changed over time after the step change. Qualitatively, the model exhibits the transient IPI overshoot relative to ISI that is observed in humans, and gradually adjusts the IPI to match the new ISI (Figure 6a, bottom). Quantitatively, the degree of overshoot in the model depends on the value of *K*. When *K* is large, the circuit is more sensitive to errors in predicting the stimulus timing and generates a larger change in *I*, resulting in more overshoot. In contrast, when *K* is small, the circuit produces less overshoot (Figure 6a, bottom).

Next, we analyzed the behavior of the model in response to a phase shift, in which a single ISI is increased or decreased, causing all subsequent stimuli to occur at a different phase without any change to the subsequent ISIs (Figure 6b, top). In response to a sudden phase shift, human subjects adjust their response times so that the initial mismatch induced by the phase shift is gradually reduced (Figure 6b, middle)^55^. When exposed to this perturbation, the model exhibited the same qualitative behavior of adjusting the outputs to gradually reduce asynchrony (Figure 6b, bottom). Quantitatively, the speed of recovery from asynchrony depends on the value of *α*, with larger values of *α* corresponding to a faster recovery (Figure 6b, bottom, circles).

Finally, we considered a perturbation in which a single stimulus is jittered temporally. This perturbation alters two consecutive ISIs in equal and opposite directions, without changing either the phase or the ISI of the subsequent stimuli (Figure 6c, top). The behavior of human subjects in response to this perturbation exhibits a characteristic biphasic error in IPI relative to ISI. For example, when a single stimulus is delayed, subjects detect the error and delay their next response accordingly. However, since the perturbation is transient, subjects have to then undo their corrective response, which they do so gradually over the course of the subsequent stimuli^56, 57^ (Figure 6c, middle). Again, the model was able to capture this response pattern qualitatively, and the dynamics of the error correction was moderately dependent on the value of *α* (Figure 6c, bottom, circles).

Together, these results demonstrate that the circuit model captures the qualitative aspects of human responses to perturbations of stimulus timing in a range of synchronization tasks. Quantitative analysis demonstrated that the dynamics of circuit motor timing depended on two key parameters of the model, *K* and *α*. These parameters give the circuit flexibility to capture the range of human behaviors observed in these tasks^52–57^. These individual differences may reflect either differential sensitivity to errors in phase and rhythmicity, or differences in internal noise.

### Circuit model implements Bayesian interval reproduction

Up to now, we demonstrated that the SAM integrates temporal information across multiple sensory inputs and allows the MPM to gradually synchronize its outputs to those inputs. We further discussed the initial inputs and update constants that will minimize the circuit’s errors in ISI estimation (Figure 3c,d). Here we offer an interpretation of the model initialization (i.e., the initial speed adopted) and adjustment (i.e., weighting given to errors) as an implementation of an optimized inference algorithm.

In the absence of any prior knowledge about the ISI of the first few beats, there is no way for the circuit (or a human) to choose an informed initial speed. Therefore, only sensory feedback can guide motor timing. However, when an observer has some prior expectation about the possible values of ISIs, the optimal strategy, as prescribed by Bayesian integration, is to integrate sensory inputs with that prior knowledge. This integration enables the system to reduce variability due to internal noise^37, 59^ and to estimate the ISI with greater precision.

Several experiments have used time interval reproduction tasks to assess whether human timing behavior is consistent with Bayesian integration^28, 60–62^. In these experiments, subjects are typically provided with a sample interval, *t*_*s*_, drawn from a fixed prior distribution, and are asked to produce an interval, *t*_*p*_, that matches *t*_*s*_ as accurately as possible. As numerous previous studies have shown^28, 60–62^, the behavior of an optimal Bayesian observer performing this task exhibits two characteristic features. First, when measurements of *t*_*s*_ are unreliable, *t*_*p*_ values are biased toward the mean of the *t*_*s*_ distribution. Second, the magnitude of these biases becomes smaller when measurements of *t*_*s*_ are more reliable.

We recently tested these characteristics in a psychophysical experiment in humans in which *t*_*s*_ was sampled from a fixed discrete uniform distribution ranging between 600 and 1000 ms^7^. To control the reliability of measurements, subjects were tested on two trial types. In one type (Figure 7a, left), which we refer to as “1-2-Go,” subjects were presented with the first two beats of an isochronous rhythm (1-2) and were asked to produce the third omitted beat (Go). In the second type (Figure 7a, right), referred to as “1-2-3-Go,” subjects were presented with the first three beats (1-2-3) and had to synchronize their response with the fourth omitted beat. We reasoned that presentation of two ISIs in the 1-2-3-Go compared to one ISI in the 1-2-Go would allow subjects to measure *t*_*s*_ more accurately and would reduce the bias. Results provided clear evidence that subjects relied on their prior knowledge since *t*_*p*_ values were biased toward the means of the *t*_*s*_ distribution (Figure 7b, purple). Moreover, biases were smaller in the 1-2-3-Go compared to the 1-2-Go condition indicating that subjects relied less on the prior when *t*_*s*_ measurement were more accurate (Supplementary Figure 2).

**Figure 7.**
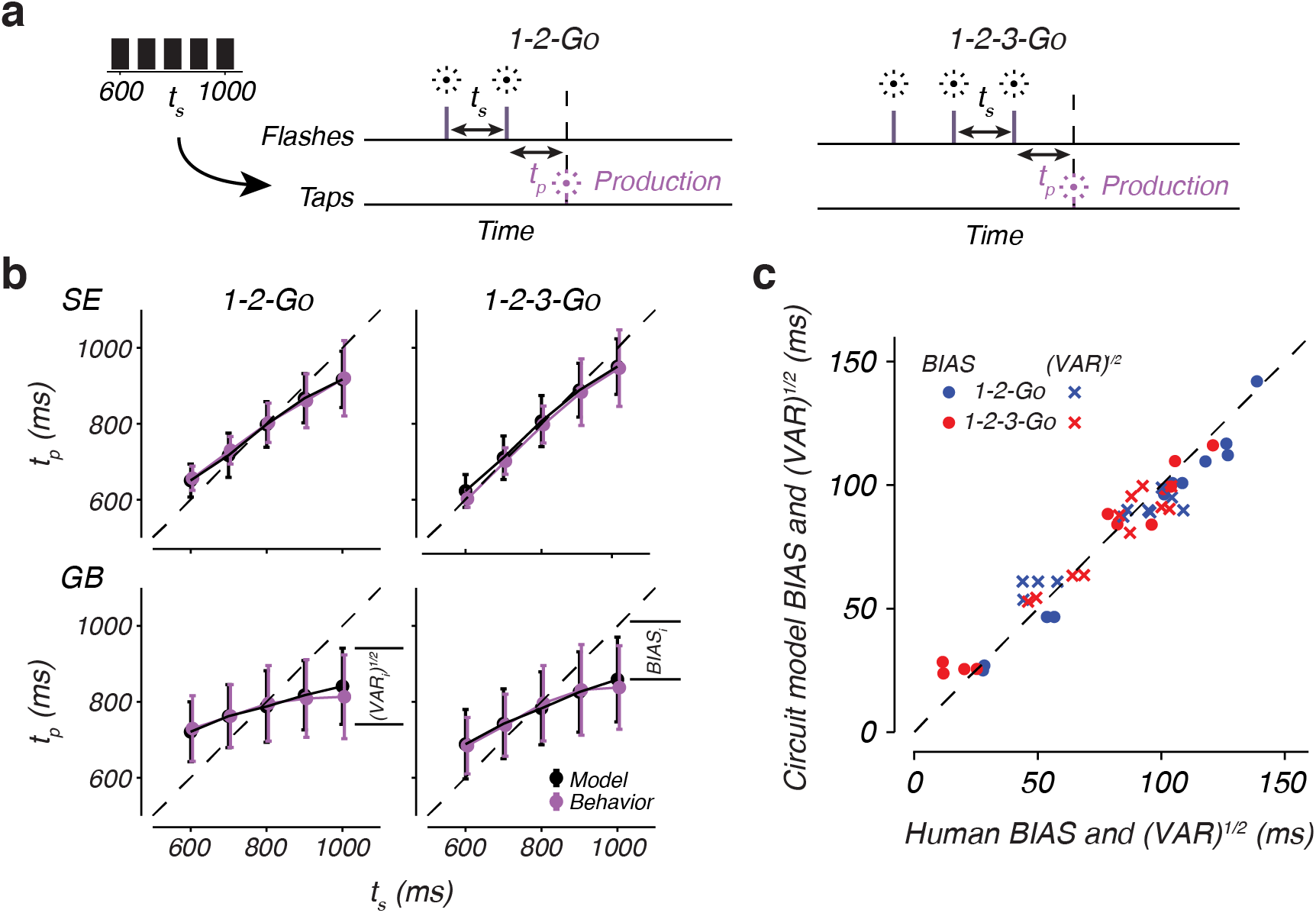
Bayesian behavior of the synchronization model. a) Schematic diagram of the Interval reproduction task. In each trial, subjects were asked to produce an interval (*t*_*p*_) that matched an interval sampled (*t*_*s*_) from a set prior distribution (left). In 1-2-Go trials (middle), subjects observed two flashes (vertical magenta lines) demarcating *t*_*s*_. In 1-2-3-Go trials, subjects observed three flashes (vertical magenta lines) demarcing *t*_*s*_ twice. b) Behavior of two example subjects (magenta) on the 1-2-Go and 1-2-3-Go tasks, together with the circuit fit to each subject (black). c) Observed BIAS and VAR of nine subjects compared to the BIAS and VAR of the circuit model fit to each subject. Circles indicates BIAS, crosses VAR, and the color indicates the trial type. Dashed line represents unity.

To test if the model could emulate characteristic features of Bayesian time interval reproduction, we simulated the model by providing the SAM with input pulses representing the beats of an isochronous rhythm (2 pulses for the 1-2-Go task and 3 for the 1-2-3-Go task). Similar to the human experiment, on each trial, the interval between pulses was drawn randomly from the sample prior distribution between 600 and 1000 ms. The behavior of the model in this task depends on three key parameters: the initial input, *I*_0_, the update constant, *K*, and the level of noise in the model, *σ*_*n*_. The parameter *I*_0_ determines the speed of dynamics before the first measurement of *t*_*s*_. This reflects the model’s initial guess about *t*_*s*_ based on prior expectations. The parameter *K* determines the strength with which the SAM updates the input based on the difference between anticipated and observed *t*_*s*_. In other words, *K* controls the sensitivity of the model to errors in anticipation. Finally, the parameter *σ*_*n*_ determines the variability of the model’s behavior due to internal noise.

Together, these parameters allow the model to capture a range of behaviors observed in human subjects. Results in Figure 7b show the behavior of the model that was fit to two human subject’s data in the 1-2-Go and 1-2-3-Go tasks (for parameter values, see Supplementary Table 1). Evidently, the model can capture several key features of Bayesian integration present in the human behavior (Figure 7b, magenta): (1) Average *t*_*p*_ increases monotonically with *t*_*s*_ (Figure 7b, black circles), (2) noise in the models causes *t*_*p*_ to vary on a trial-by-trial basis for the same *t*_*s*_ (Figure 7b, black error bars), (3) average *t*_*p*_ is biased toward the mean of *t*_*s*_ distribution (Figure 7b, deviation from the dashed unity line), and (4) the magnitude of bias is smaller in the 1-2-3-Go compared to the 1-2-Go trials (*z* = 2.7, *p <* 0.01, one-sided Wilcoxon signed rank test; Supplementary Figure 2). Moreover, the speed of dynamics before sensory feedback corresponded to an ISI of 850.25 ± 39.26 ms (see Supplementary Table 1), consistent with anticipating the ISI based on an approximation of the prior mean.

To further compare the model’s behavior to that of the human subjects, we partitioned the root-mean-square error (RMSE) between *t*_*p*_ and *t*_*s*_ to two terms, a BIAS terms that measures the overall bias, and a VAR term that measures average variability across all values of *t*_*s*_ (see Methods). The model was able to capture the observed BIAS and VAR across subjects in both the 1-2-Go and 1-2-3-Go (Figure 7c), indicating that it can correctly implement the bias-variance tradeoff exhibited by humans during interval reproduction. In this model, the update constant plays a central role in determining the bias-variance tradeoff. Larger values of *K* lead to less bias and vice versa (compare Figure 7b, top and bottom). For a Bayesian observer, the magnitude of bias is determined by noise in the measurements of *t*_*s*_ and by the imposed cost function. Accordingly, the value of *K* in the model was adjusted to achieve a level of bias-variance tradeoff that is inversely related with the inherent noise in the model (*σ*_*n*_) and the operative cost function (*r* = *−*0.94, *p* = 0.0002; see Supplementary Table 1).

### Bayesian synchronization/continuation

As our final test, we examined the model in the context of a task that combines all the previous capabilities including sensory anticipation, motor timing and Bayesian integration, which we refer to as the Bayesian synchronization/continuation task. The basic synchronization/continuation task requires subjects to tap synchronously with a metronome (synchronization phase) and continue to tap with the same tempo without the metronome until instructed to stop (continuation phase). In the classic version of this task, subjects have no uncertainty about the ISI of the metronome as it is kept constant across consecutive trials. Recently, humans were tested on a variant of this task in which ISI for each trial was sampled randomly from a prior distribution^38^. While performing this task, humans exhibit distinct patterns of biases in their IPIs. During synchronization, responses were biased toward the mean of the ISI distribution and the magnitude of the bias decreased with the number of synchronization stimuli (Figure 8a, purple lines). These observations are consistent with the Bayesian integration scheme that we discussed previously in the context of a time interval reproduction task (Figure 7). During the continuation phase, the bias persisted and its magnitude increased substantially (Figure 8a, red lines) for all subjects (Figure 8b). This increase in bias was manifested as a drop of sensitivity of IPI to ISI (shallower slope relating IPI to ISI) as well as an overall bias of IPIs toward shorter ISIs (Figure 8c, left).

**Figure 8.**
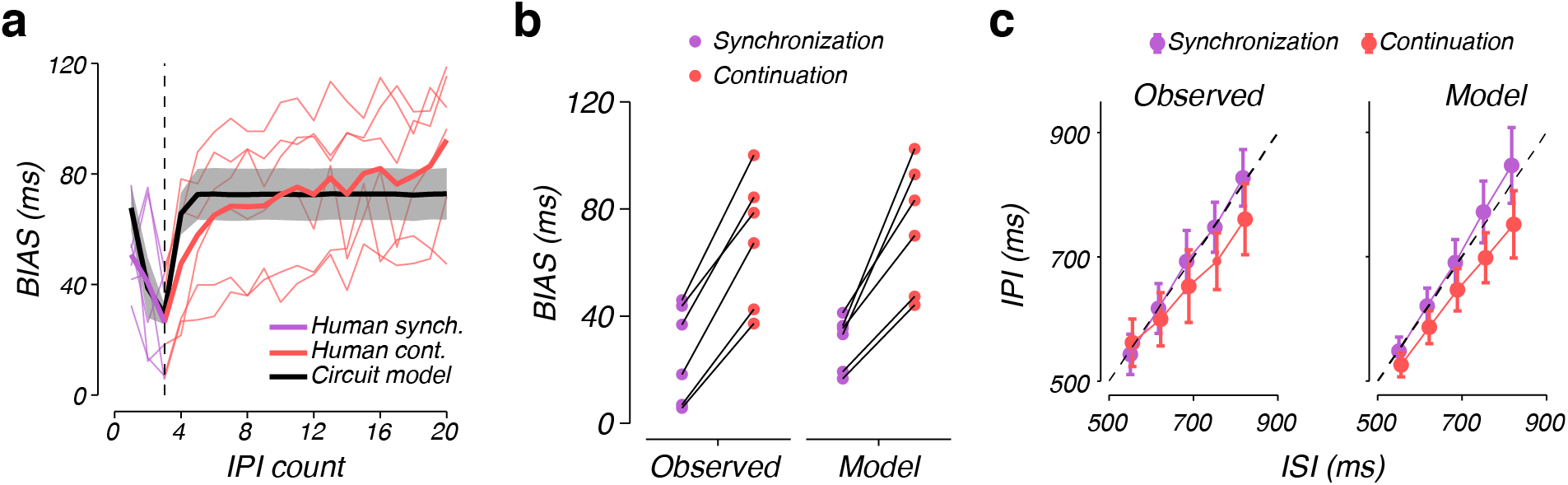
Circuit model captures systematic biases in human behavior during Bayesian synchronization/continuation. a) Overall BIAS in the synchronization/continuation task. BIAS was calculated as in Figure 7. Purple and red lines indicate BIAS for human subjects during synchronization and continuation, respectively. Thin lines indicate individual subjects. Thick lines indicate the mean across subjects. The black line with shading indicates the mean and standard deviation of the BIAS in the circuit model fit to individual subjects. The vertical dashed line marks the transition from synchronization to continuation. b) Observed and model BIAS for individual subjects during each phase of the task. Lines connect data points for each subject or their respective model fits. c) Inter-production interval (IPI) for different inter-stimulus intervals (ISI) for an example subject (left) and the circuit model fit to that subject’s behavior (right). Dashed lines indicate unity.

To test whether the circuit model could capture this nontrivial pattern of biases during the synchronization/continuation task, we fitted the model (*I*_0_, *K*, and *α* parameters) to individual subject’s behavior (see Methods; Supplementary Table 2). The fitted models were able to capture the pattern of biases: IPIs were biased towards the mean ISI, and the magnitude of the bias decreased during synchronization and increased during continuation (Figure 8a, black line). Like humans, the increase in bias during continuation was due to a combination of decreased sensitivity to the ISI (decrease in slope of IPI-ISI relationship, *z* = 2.1, *p* = 0.02, one-sided Wilcoxon signed rank test) and an overall shift towards shorter IPIs (decrease in IPI of the middle interval, *z* = 2.1, *p* = 0.02, one-sided Wilcoxon signed rank test; Figure 8c, right; Supplementary Figure 3).

In the model, this pattern of biases is explained by the augmented input, ∆*I*, to the MPM. Following the final stimulus input, the SAM no longer resets and instead, it proceeds toward the terminal fixed point. As a result, its output, *y*_*s*_, becomes fixed at a value greater than *y*_0_. However, *y*_*p*_ continues to oscillate between values smaller than *y*_0_ because it is reset by the motor output. This mismatch makes ∆*I* negative and decreases the overall input to the MPM. The decreased input, in turn, speeds up the dynamics of the MPM and shortens IPIs during the continuation phase across all ISIs. Further, the impact of this negative ∆*I* is integrated over time, leading to an asymmetric impact on the speed of MPM dynamics associated with a short ISI compared to a long ISI. Specifically, longer ISIs lead to a greater increase in speed than shorter ISIs, accounting for the decreased sensitivity to the ISI during continuation. It is important to note that these results require that the SAM does not reset. Therefore, an alternate circuit configuration in which the output of the MPM would reset the SAM would not be able to capture the increase in biased during continuation. This finding validates our assumption that resetting the SAM relies on sensory inputs and not motor outputs.

## Discussion

To successfully control motor outputs in time, neural circuits must integrate contextual information with sensory input to control the dynamic evolution of neural activity^20, 23, 63–65^. The implementation of effective control, however, is challenging in the presence of uncertainty and time delays between motor output and the collection of sensory information^30^. Our results demonstrate that pairing a module that executes actions with an internal simulation of that module allows a simple circuit mechanism to achieve closed loop control over timing behavior. Adopting this control architecture allows the circuit to generate dynamic activity that anticipates upcoming sensory input, effectively bridging the time delay between action initiation and subsequent feedback. By overcoming this delay, the sensory anticipation module creates a basis for adjusting the subsequent dynamics based on feedback without overt motor output. These adjustments to the control of internal processes can then be applied to the circuits responsible for motor output, allowing the appropriate selection of dynamics to achieve the desired action timing.

The simulator enables the integration of feedback following long time delays and allows the circuit to achieve synchronization in a manner consistent with humans^43, 44, 50, 51, 54^. By using the output of the simulator in combination with sensory input, the system can update input controlling the speed of dynamics. The interaction between the initial conditions of the circuit (e.g. *I*_0_) and the updating mechanism results in systematic biases observed during human interval reproduction behavior^7, 28, 61, 62^. Because *I*_0_ effectively sets motor timing without sensory feedback, this input represents prior knowledge of the mean of the possible desired intervals. This prior knowledge, when combined with the updates through errors weighted by the parameter *K* to integrate sensory feedback, results in action timing consistent with Bayesian interval reproduction. Importantly, the system captures non-trivial features of human behavior during Bayesian synchronization/continuation. The interaction of the predictive mechanism with the phase-correction mechanism generates the decrease in bias during synchronization and increase in bias during continuation, consistent with the pattern of biases in humans performing an identical task^38^.

Previous modeling efforts focusing on the psychophysics of interval estimation have proposed iterative error correction algorithms to coordinate motor timing with sensory timing^7, 40, 43, 44, 51^. We complement this work from human psychophysics by implementing these algorithms as a circuit model that generates predictions and error signals without an overt movement. We base our circuit on a core module that implements speed control in a manner consistent with neural responses^20^. Pairing a motor production module with a sensory anticipation module allows the circuit to generate internal activity that predicts the onset of the next stimulus and uses this prediction to compute an error signal that serves to adjust subsequent timing behavior. Critically, the simulator generates anticipatory signals related to the timing of sensory feedback in the absence of overt motor output. The circuit model developed here could, therefore, serve as a conceptual basis for a mechanistic understanding of the predictive processes thought to be implemented by humans during perceptual timing tasks^40, 66, 67^.

Previous circuit models of timing largely fall into three categories, each with their own strengths and weaknesses. The first class of models is based on the accumulation of ticks of a central clock^68–71^. Similar to our circuit model, these clock-accumulator models generate ramping activity often observed in subsets of single units during neurophysiological experiments^21, 22, 72^. However, these models fail to explain the diversity of dynamic responses in populations of neurons thought to contribute to timing^20, 23, 65, 73–75^. The second category of models is based on dynamical systems. These models rely on the rich set of dynamics generated by neurons connected in a local circuit to represent the passage of time^9, 20, 76–78^. These models generate complex dynamics that are strikingly similar to those observed in local populations, but it is unclear whether and how they can flexibly integrate sensory and motor feedback. The third category of model commonly used to capture sensorimotor timing are circuits that rely on oscillatory modules^45–47, 79–81^. These models implement coupling between many oscillatory modules in a circuit to achieve sophisticated timing behaviors that integrate sensory information across a range of timescales^45^. While these models can capture complex timing behaviors, it is not clear how they relate to observed responses from individual neurons in the brain regions causally involved in timing^20^. Our simple circuit model provides an understanding of the link between these previous models. First, it establishes a temporal control mechanism based on ramping activity, which is observed in individual neurons^22, 23, 26, 82, 83^. However, the ramping in the model results from circuit-level recurrent dynamics that, in principle, can be extended to larger circuits capable of generating more complex dynamics^9, 20, 84^. Second, the model generates oscillations at its output. However, unlike more abstract models comprised of inherently oscillatory units, our model establishes oscillations through dynamic interactions between anticipatory and feedback mechanisms responsible for speed control that have been characterized in-vivo and in larger recurrent circuit models^9, 20, 85^.

Finally, the wiring of our circuit model might provide insight into the individual functional contributions of the cortical and subcortical systems important to action and perceptual timing^86^. One node, the basal ganglia, has been heavily implicated in action timing through lesion studies^87^ and physiological evidence^20, 65, 88, 89^. An important principle of the basal ganglia function is the inhibition of downstream neural activity^90^. Given their proposed function as a competitive selection mechanism^91^, these inhibitory pathways may be the substrate for implementing the mutual inhibitory interactions needed for the temporal control of movements. Another key node, the cerebellum, has also been linked to timing through lesion studies^92^ and physiology^74, 93^, and modeling^38^. A hallmark of cerebellar function is the detection and correction of sensory errors during sensorimotor control^94, 95^, which is the key function of the sensory anticipation module in our model. Finally, the output of both the basal ganglia and cerebellum are sent trans-thalamically to regions of the frontal cortex involved in sensory and motor timing^96–98^. Accordingly, we speculate that the integration of the sensory anticipation and motor production modules in our model may rely on frontal cortical areas.^97^.

## Methods

### Basic circuit module (BCM) for interval production

The fundamental circuit architecture consists of four rate units, *I*, *u*, *v*, and *y*, which are governed by the following set of equations

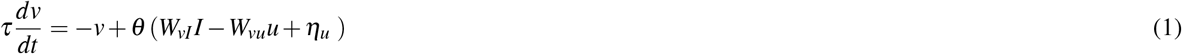

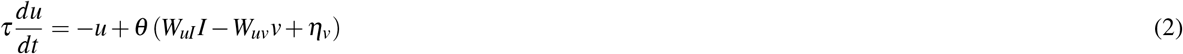

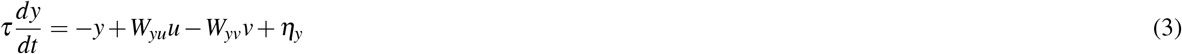

Where *τ*, the time constant of each unit, was set to 100 ms and *θ* (*x*), the activation function of each unit, was specified by *θ* (*x*) = 1*/*[1 + exp(− *x*)]. *W*_*uI*_, *W*_*uv*_, *W*_*vu*_, and *W*_*vI*_ specify the weighting of the interactions between units and each was set to 6 for all simulations. Similarly, we set *W*_*yu*_ = *W*_*yv*_ = 1. Noise in the system is modeled by zero mean Gaussian white noise inputs, represented by the variables *η*_*u*_, *η*_*v*_, and *η*_*y*_ for unit *u*, *v*, and *y*, respectively. The units were initialized at *u* = 0.7, *v* = 0.2, *y* = 0.5, *I* = *I*_0_ where *I*_0_ is a free parameter. All simulations were carried out using Euler’s method with a step size of 10 ms. Noise variables were independently sampled at every time point from a Gaussian distribution with mean zero and standard deviation specified by *σ*_*n*_.

### Motor planning module (MPM)

The motor planning module (MPM) consists of four units *I*, *u*_*p*_, *v*_*p*_, and *y*_*p*_, which are simulated based on the following equations.

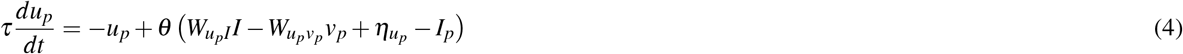

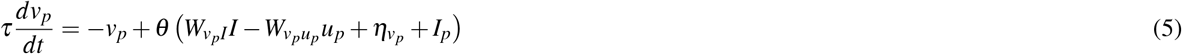

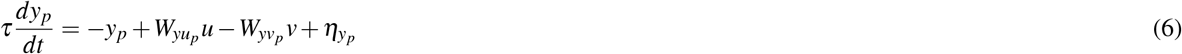

*I*_*p*_ specifies a transient input to *u*_*p*_ and *v*_*p*_ that serves to reset the system when *y_p_ > y*_0_. We set *I*_*p*_ to 50 and *y*_0_ to 0.7 for all simulations. Noise, represented by the variables *η*_*up*_, *η*_*vp*_, and *η*_*yp*_, was injected into each unit as in the BCM. All other parameters are the same as the BCM.

### Sensory anticipation module (SAM)

The sensory anticipation module consists of four units *I*, *u*_*s*_, *v*_*s*_, and *y*_*s*_. The system evolves according to the equations

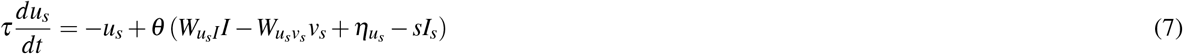

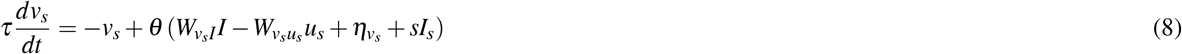

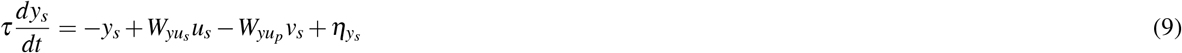

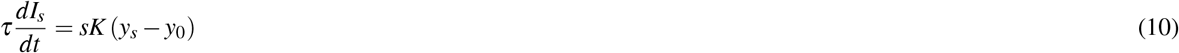

where *K* is a free parameter and *η* indicates noise input to each unit. The module also receives a binary input *sI*_*s*_, where *s* represents the visual stimulus (*s* = 1 when the stimulus is on and *s* = 0 when the stimulus is off) and *I*_*s*_ is set to 50. This input serves to reset the values of *u*_*s*_ and *v*_*s*_ at the time of each stimulus. All other parameters are the same as the BCM.

Sequences of stimuli were implemented by setting *s* = 1 periodically. Each stimulus presentation lasts 10 ms. The units were initialized at *u* = 0.7, *v* = 0.2, *y* = 0.5, and *I* = *I*_0_ where *I*_0_ is a free parameter. The dynamics were allowed to run forward for 750 ms before the first stimulus was presented. This initialization period serves to put the circuit closer to steady-state so that the effect of reset signals is consistent across different ISIs. *I* does not update during the first presentation of the stimulus (by effectively setting *K* = 0 during the first stimulus presentation).

Optimization of SAM parameters was achieved by simulating the module with 100 different pairs of *I*_0_ and *K* for each *σ*_*n*_. *K* was uniformly sampled from values between 1 and 8.0, and *I*_0_ was sampled from values between 0.77 and 0.79. *N* stimuli were presented to the model where *N* = 2, 3, …, 10.

For each set (*σ*_*n*_, *I*_0_, *K*, *N*), the model was run as described in Circuit model behavior and the RMSE was calculated

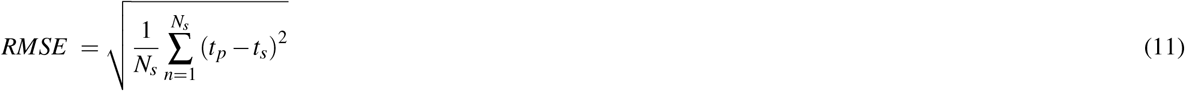

where *N*_*s*_ = 500 is the total number of iterations across all *t*_*s*_. The pair (*I*_0_, *K*) that results in the smallest RMSE was recorded. The procedure was repeated 10 times for each *σ*_*n*_ to obtain a distribution of the optimal (*I*_0_, *K*).

In figure 3c, *I*_0_ and *K* were picked from a 30 *×* 30 grid which spans the same intervals, and *N*_*s*_ = 5000.

### Behavioral Tasks

We used a suite of behavioral tasks to test circuit model timing performance. In all tasks, we performed numerical simulations of the dynamical equations expressed above using Euler’s method and a time step of 10 ms. All behavioral experiments were performed with the approval of the Committee on the Use of Humans as Experimental Subjects at MIT after receiving informed consent.

#### Periodic production

To achieve periodic production, we simulated the motor planning module in isolation at four different levels of input, *I*, uniformly spaced from 0.75 to 0.78, for 40 s at each level. To measure the performance of the model, we calculated the IPI as

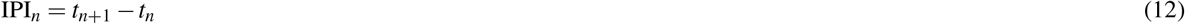

Where *t*_*n*_ was the time of the *n*^th^ action produced by the model. For all levels of input we set *σ*_*n*_ = 0.01. To compare the timing behavior of the model across levels of input, we calculated the mean and standard deviation of the first 40 IPIs generated by the circuit.

#### ISI tracking

Circuit model synchronization was tested in a randomized ISI tracking task. The ISI was defined as

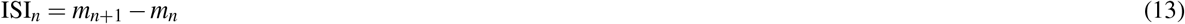

Where *m*_*n*_ was the time of the *n*^th^ stimulus. Each trial of the task, we initialized the ISI at a value of 800 ms and simulated a sensory input with 20 consecutive 800 ms intervals. After the first block of 20 ISIs, the ISI for the next 20 intervals was selected at random a discrete uniform distribution between 600 and 900 ms with four possible values. This process was repeated four total times, to generate a sequence of 100 total intervals.

Circuit parameters were selected as follows: *I*_0_ was set to 0.771; *K* was selected to be 2; *α* was chosen to be 0 for simulations with no augmented input (e.g. Figure 4) and 0.1 for simulations with augmented input (e.g. Figure 5). *I*_0_ was set such that the MPM generates an 800 ms IPI on average, matching the initial ISI of every trial. This ensured that differences in circuit output resulted from updates to the value of *I*, rather than the value of *I*_0_ selected.

The *n*^th^ IPI was quantified as in equation 12 and compared to the *n*^th^ ISI. The asynchrony between an action and the stimulus presented at time *m*_*n*_ was defined as

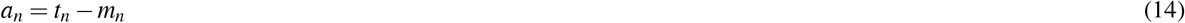

Where *t*_*n*_ was the time of the action which was closest in time to *m*_*n*_. The relative phase of the action relative to the *n*^th^ stimulus, *φ*_*n*_, was calculated as

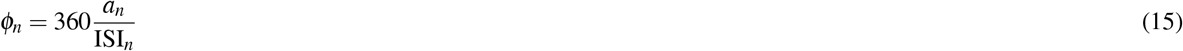

Where the phase is in units of degrees. For all simulations, we set *σ*_*n*_ = 0.01.

#### ISI perturbations

The basis of the mechanisms behind human synchronization behavior are generally probed using perturbations of the ISI after an experimental subject has reached steady-state synchronization performance to a given ISI (see^50^ for a review). To compare the circuit model synchronization performance to that of humans, we explored the behavior of the model in response to three common ISI perturbations: (1) A *step change* in the ISI, where the circuit synchronized to a stimulus with an ISI of 800 ms before the ISI was stepped to 1000 ms for all subsequent ISI. (2) A *phase shift* in the stimulus timing, where the circuit synchronized to a stimulus with a 500 ms ISI before the timing of stimuli was shifted by increasing a single ISI to 600 ms. All subsequent ISIs were 500 ms. (3) *Stimulus jitter*, where the circuit synchronized to an ISI of 500 before the timing of a single stimulus is perturbed, while subsequent stimuli remain in phase with the stimulus before the perturbation. To accomplish this, we perturbed two successive ISIs; the first was increased to 600 ms and second was decreased to 400 ms.

We simulated 1000 trials of each perturbation type and calculated the asynchrony associated with each action as in equation 14 and the mean IPI between actions as in equation 12. To ensure the circuit model was fully synchronized before a perturbation, we simulated 30 ISIs of the same duration before applying the perturbation. The circuit model was simulated with *I*_0_ = 0.771 and *σ*_*n*_ = 0.005. The values of *K* and *α* were varied to explore the behavior of the model with different sensitivities to errors in simulation and synchronization.

#### 1-2-Go and 1-2-3-Go

We compared the circuit model interval reproduction behavior to that of humans performing a timing task we refer to as 1-2-Go and 1-2-3-Go. The methods used for testing human interval reproduction are summarized briefly here. Please see our behavioral paper for a full description of the task and methods^7^.

##### Subjects and apparatus

The behavior of nine human subjects was analyzed. Subjects sat in a dark, quiet room at a distance of approximately 50 cm from a display monitor. The display monitor had a refresh rate of 60 Hz, a resolution of 1920 by 1200, and was controlled by a custom software (MWorks; http://mworks-project.org/) on an Apple Macintosh platform.

##### Interval reproduction task

The task consisted of two randomly interleaved trial types “1-2-Go” and “1-2-3-Go”. On 1-2-Go trials, two flashes demarcated a sample interval (*t*_*s*_). On 1-2-3-Go trials, three flashes demarcated *t*_*s*_ twice. For both trial types, subjects had to reproduce *t*_*s*_ immediately after the last flash by pressing a button on a standard Apple keyboard. On all trials, subjects had to initiate their response proactively and without any additional cue. Subjects received graded feedback on their accuracy.

Each trial began with the presentation of a 0.5°circular fixation point at the center of a monitor display. The color of the fixation cued the trial type. After a 2 second delay, a warning stimulus and a trial cue were presented. The warning stimulus was a white circle that subtended 1.5°and was presented 10°to the left of the fixation point. The trial cue consisted of 2 or 3 small rectangles 0.6°above the fixation point (subtending 0.2° × 0.4°, 0.5°apart) for the 1-2-Go and 1-2-3-Go trials, respectively. After a random delay with a uniform hazard (250 ms minimum plus an interval drawn from an exponential distribution with mean of 500 ms), flashes demarcating *t*_*s*_ were presented. Each flash lasted 6 frames (100 ms) and was presented as an annulus around the fixation point with an inside and outside diameter of 2.5°and 3°, respectively. *t*_*s*_ was sampled from a discrete uniform distribution ranging between 600 and 1000 ms with a 5 samples. To help subjects track the progression of events throughout the trial, after each flash, one rectangle from the trial cue would disappear (starting from the rightmost). The produced interval (*t*_*p*_) was measured as the interval between the time of the last flash and the time when the subject pressed the response key. We discarded any trial when the subject responded before the second (for 1-2-Go) or third (for 1-2-3-Go) flash and any trial where the response was 1000 ms after the veridical *t*_*s*_. We further used a model based approach to identify “lapse trials,” or trials in which *t*_*p*_ was not related *t*_*s*_^7^. We then calculated the mean *t*_*p*_ conditioned on *t*_*s*_ and trial type.

##### Circuit model behavior

The sensory anticipation module was run for 100 simulated trials for each *t*_*s*_ (600 ms, 700 ms, 800 ms 900 ms, 1000 ms). To simulate the 1-2-Go task, the SAM was presented with 2 stimuli that are separated by the selected *t*_*s*_. After the second stimuli was presented, the first time *T* for which *y_s_ > y*_0_ was defined as the production time of the model, and *t*_*p*_ is the difference between *T* and the time of the second stimulus. The 1-2-3-Go task was simulated similarly but with 3 stimuli.

##### Model fitting

For each subject, we calculated the mean and standard deviation of *t*_*p*_ for each *t*_*s*_ in both the 1-2-Go and 1-2-3-Go tasks (10 task conditions in total). We call these values *µ*_1,subject_, *ST D*_1,subject_, …, *µ*_10,subject_, *ST D*_10,subject_ (one pair of values for each condition).

For each set of parameters (*σ*_*n*_, *I*_0_, *K*), the model was run as described in Circuit model behavior. We calculated the mean and standard deviation of the model’s *t*_*p*_ for each *t*_*s*_, and for both the 1-2-Go and 1-2-3-Go tasks. We call these values *µ*_1,model_, *STD*_1,model_,…, *µ*_10,model_, *ST D*_10,model_.

The fitting was done by alternating between fitting *σ*_*n*_ and jointly fitting (*I*_0_, *K*). The parameters were uniformly and independently sampled from the following intervals: [1.0, 8.0] for *K*, [0.77, 0.79] for *I*_0_ and [0.005, 0.4] for *σ*_*n*_. For each step, 100 sets of parameters were sampled and the optimal set was used to update the parameters.

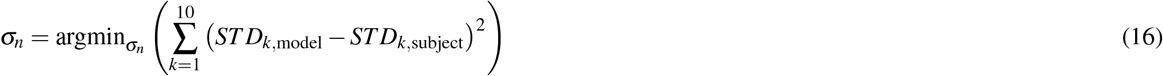

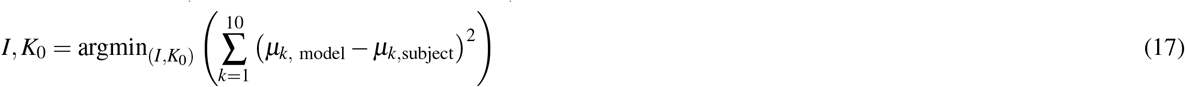

To evaluate the anticipated *t*_*s*_ associated with *I*_0_ and *σ*_*n*_, we initialized the model with *I*_0_ and presented a single stimulus to the model 750 ms after initialization. The anticipated *t*_*s*_ was the production time of the model after the stimulus is presented.

#### Synchronization/continuation

We compared the circuit model to the behavior of humans performing a synchronization/continuation task in which the interval between flashes (ISI) was selected at random each trial from a set distribution. The methods used for testing human experiments are summarized briefly here. Please see Narain et. al. (2018) for a full description of the task and methods^38^.

##### Stimulus and apparatus

We analyzed the behavior of 6 human participants. Stimuli were viewed from a distance of approximately 67 cm in a dark, quiet room. The display monitor had a refresh rate of 60 Hz, a resolution of 1920 by 1200, and was controlled by a custom software (MWorks; http://mworks-project.org/) on an Apple Macintosh platform.

##### Synchronization/continuation task

Each trial began with the presentation of a red circular fixation stimulus (diameter = 0.75 degrees visual angle) in the center of the screen. After a variable delay (200 ms plus an additional amount of time which was exponentially distributed with mean = 300 ms and a maximum value of 2300 ms), a synchronization stimulus was flashed four times with an interstimulus interval (ISI) chosen from a discrete uniform distribution (five intervals, minimum = 550 ms, maximum = 817 ms). The flashing stimulus was a gray annulus centered around fixation with an inner diameter of 1 degree and outer diameter of 1.25 degrees. Participants were instructed to tap a response key synchronously with the third and fourth synchronization flashes and continue tapping to reflect the ISI until instructed to stop. The number of continuation taps required was three plus an exponentially distributed number with a mean of nine and a maximum of 22. The first interproduction interval (IPI) was defined as the interval between the middle of the second flash and the first key press. Subsequent (IPIs) were defined as the interval between successive key presses.

Biases in IPIs were calculated according to the following procedure. For each interval, ISI_*i*_, we found the mean IPI for the *k*^th^ interval in the sequence 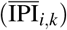. The mean squared bias for each *k* was defined as

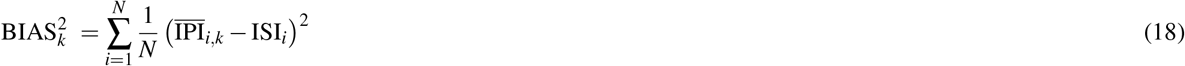

where *N* = 5 is the number of target intervals.

##### Circuit model behavior

We simulated 21 trials of the full synchronization model for each ISI. In each trial, the stimuli presented to the model were a series of three flashes separated by the ISI of that trial. After the three flashes, the stimulus was set to *s* = 0 and the model was run and allowed to produce 17 more productions. The IPI of the circuit was defined as the time difference between successive productions.

##### Model fitting

We fixed *σ*_*n*_ at 0.01 and varied three free parameters *K*, *I*_0_, and *α*. For each subject, we simulated 100 different parameter combinations with *K* selected from the interval [0.01, 5], *I*_0_ from the interval [0.76, 0.78] and *α* from the interval [0.01, 0.1]. For each combination of parameters, the model was run as described in *Circuit model behavior* and bias was calculated as in equation 18. We then found the combination of parameters that minimized the mean squared errors between the BIAS_*k*_ observed in the subject and the BIAS_*k*_ of the circuit, across *k* = 1,…, 20.

Synchronization bias was quantified as the bias for *k* = 3, and continuation bias was the bias for *k* = 7.

## Acknowledgements

The authors thank Jing Wang, Morteza Saraf, Nicolas Meirhaeghe, and Alexandra Ferguson for their insightful comments on this manuscript. M.J. is supported by NIH (NINDS-NS078127), the Sloan Foundation, the Klingenstein Foundation, the Simons Foundation, the McKnight Foundation, the Center for Sensorimotor Neural Engineering, and the McGovern Institute.

## Author contributions

SWE conceived of the extension of the basic circuit module. SWE developed, implemented and characterized the motor planning module. NML developed, implemented and characterized the sensory anticipation module. SWE developed the circuit for synchronization task and characterized model performance. NML compared model performance to humans in the 1-2-Go, 1-2-3-Go, and synchronization/continuation tasks. MJ supervised the project. All authors contributed to the interpretation of results and writing of the manuscript.

## Supplementary Materials

**Supplementary Figure 1.**
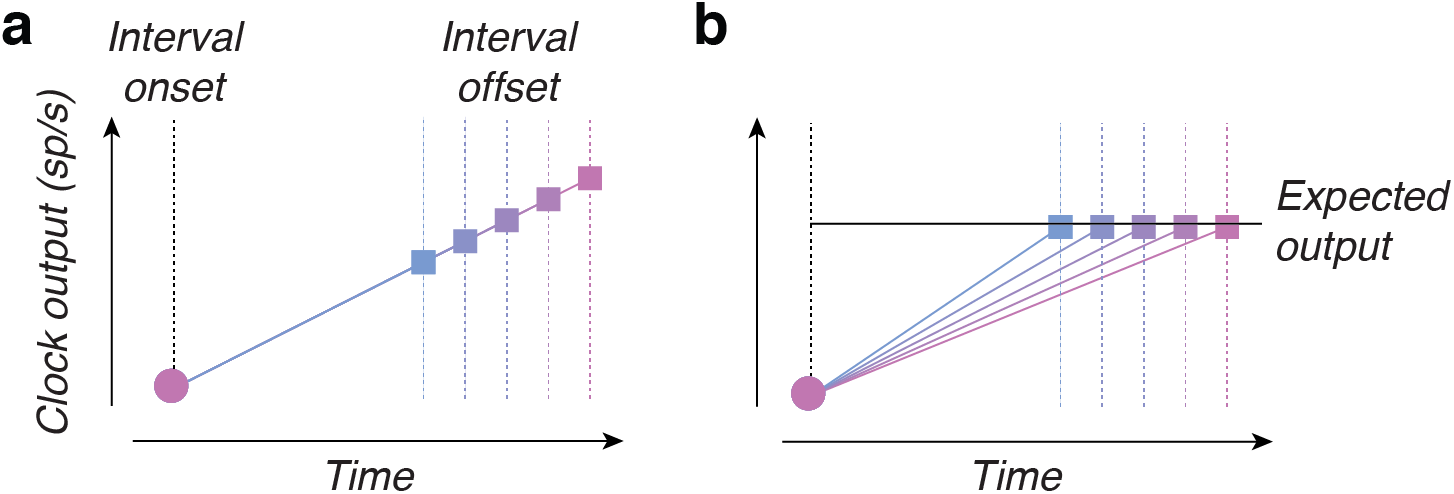
Absolute and predictive timing mechanisms. a) Absolute timing mechanism. In this mechanism, neural systems integrate the ticks from a central clock, generating activity that increases monotonically (colored solid lines) from interval onset (black dashed line) to interval offset (colored dashed lines). Because the rate of increase does not depend on the interval, the output serves as a continuous estimate of the elapsed time, and the level of activity at the time of the interval offset (squares) signals interval duration; low output activity would indicate a short interval (blue square) and high output activity would indicate a long interval (pink square). b) Predictive timing mechanism. In this mechanism, the rate of increase is adjusted proactively such that activity increases quickly in anticipation of a short interval (solid blue line) and slowly in anticipation of a long interval (solid pink line). The output of the timer now provides a continuous readout of time relative to the anticipated interval duration. If the rate of increase is set correctly, the output at the time of interval offset (squares) will match the expected output (black horizontal line). Output that is less than the expectation indicates the interval is shorter than anticipated while output that is larger than expectation indicates the interval is longer than anticipated.

**Supplementary Figure 2.**
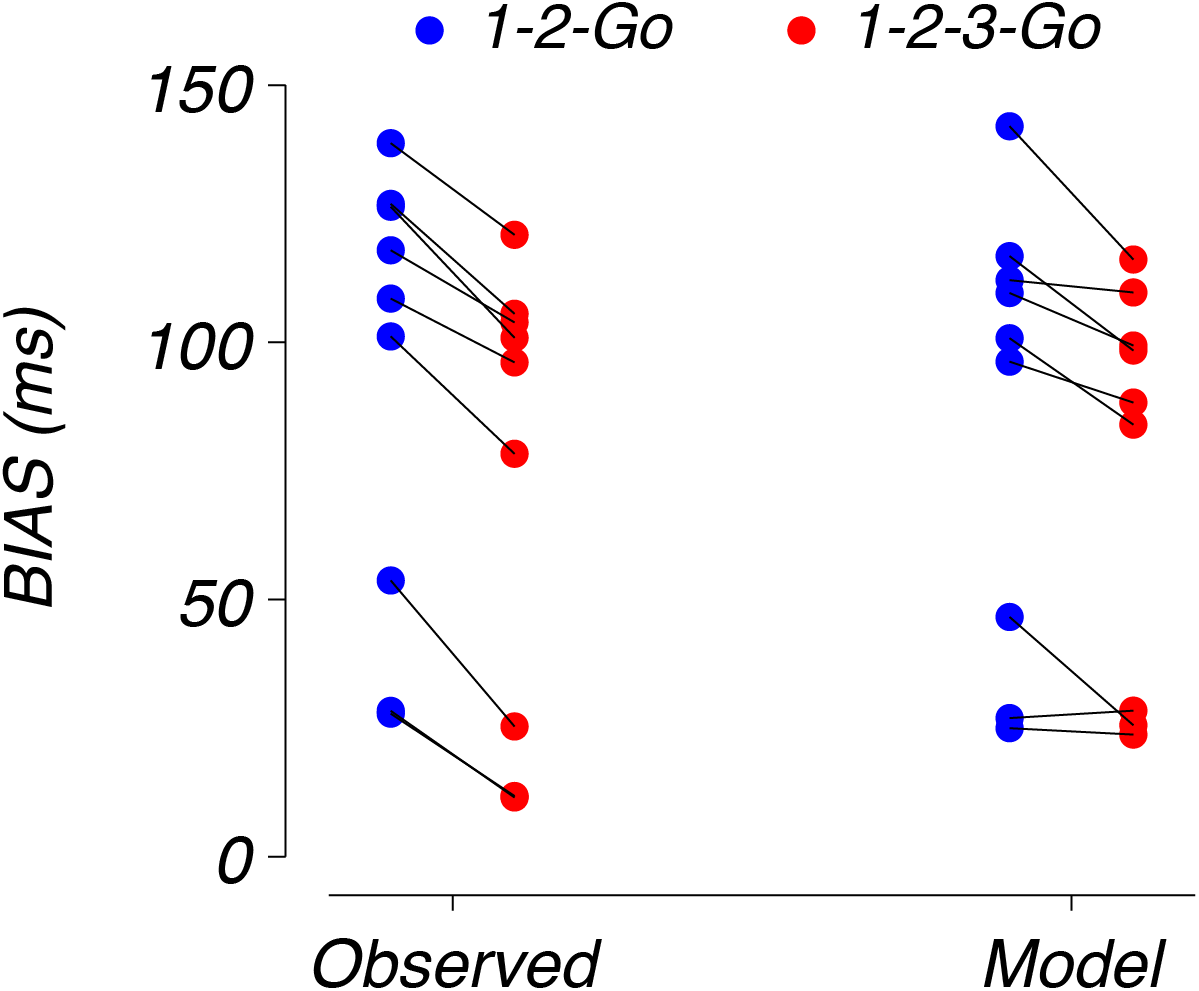
Subject and model BIAS in 1-2-Go and 1-2-3-Go. Left: observed subject BIAS (see Methods) in 1-2-Go and 1-2-3-Go. Right: BIAS of the model in 1-2-Go and 1-2-3-Go. Dots represent the data for each condition and lines connect data points across conditions for the same subject.

**Supplementary Figure 3.**
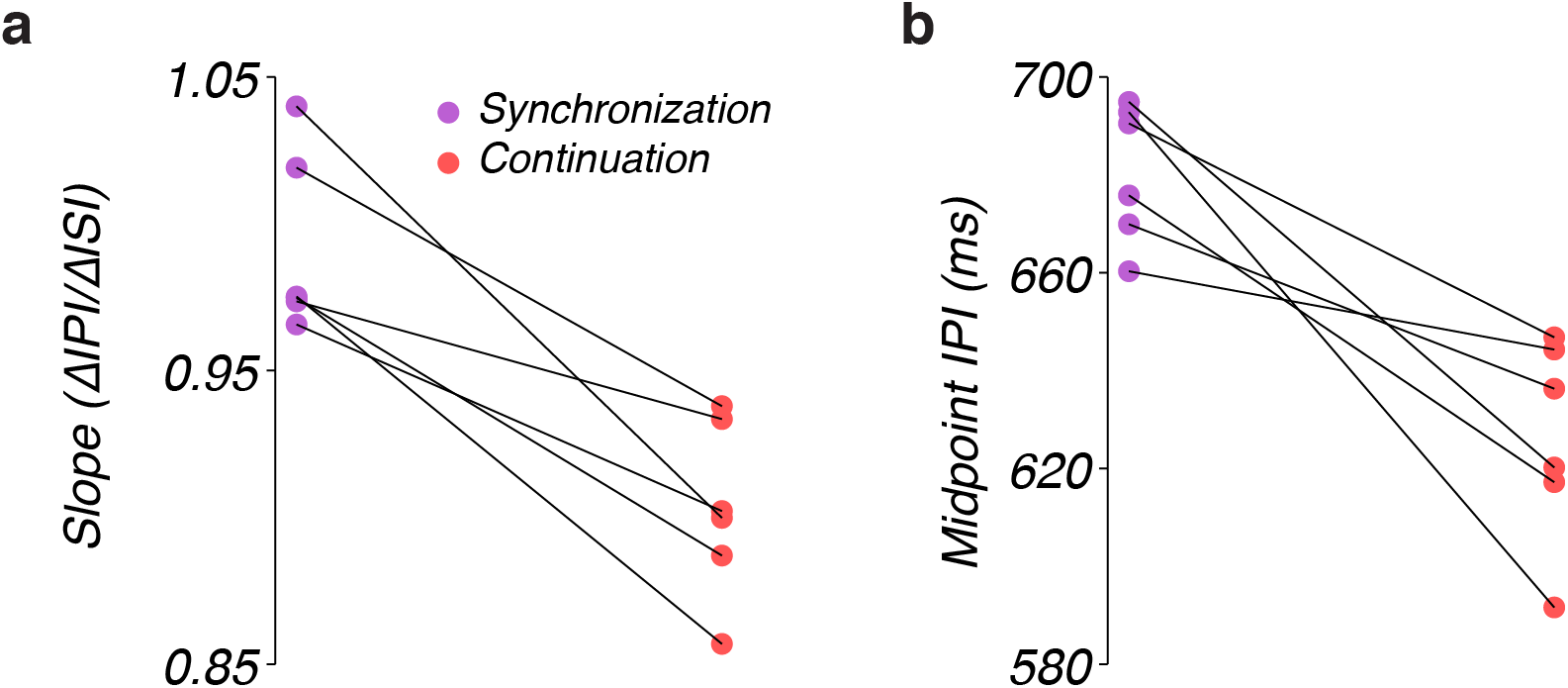
Sensitivity of the interproduction interval (IPI) of the circuit model to interstimulus interval (ISI) and the overall offset of the IPI during synchronization and continuation. a) Slope of the regression line fit to IPI versus ISI data during synchronization and continuation. Data points plot the slope of individual subjects and lines connect synchronization and continuation data for each subject. b) Mean IPI during synchronization and continuation in response to an ISI of 700 ms. The 700 ms ISI was the midpoint of the prior distribution of ISIs. Therefore, the mean IPI in response to the 700 ms ISI indicates the degree to which the overall IPI response curve is shifted relative to the midpoint of the data.

**Supplementary Table 1.**
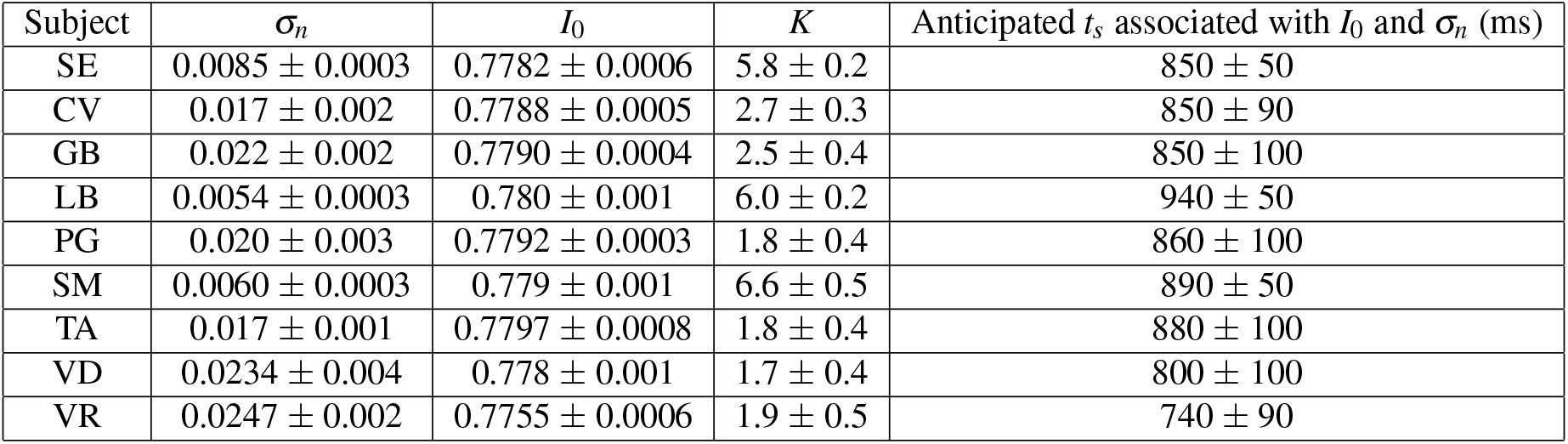
Parameters of model fits to the 1-2-Go and 1-2-3-Go behavior (mean ± standard deviation across 5 optimization runs for each subject).

**Supplementary Table 2.** Parameters of model fits to the synchronization/continuation behavior (mean ± standard deviation across 20 optimization runs for each subject).

| Subject | $I_0$ | $K$ | $\alpha$ |
| --- | --- | --- | --- |
| AL | $0.767 \pm 0.002$ | $1.9 \pm 0.6$ | $0.08 \pm 0.01$ |
| ER | $0.765 \pm 0.003$ | $4.2 \pm 0.6$ | $0.077 \pm 0.009$ |
| FK | $0.771 \pm 0.002$ | $4.0 \pm 0.6$ | $0.05 \pm 0.01$ |
| KL | $0.768 \pm 0.002$ | $1.7 \pm 0.6$ | $0.08 \pm 0.01$ |
| MW | $0.770 \pm 0.003$ | $4.4 \pm 0.4$ | $0.055 \pm 0.008$ |
| RC | $0.766 \pm 0.002$ | $3.1 \pm 0.7$ | $0.09 \pm 0.01$ |

